# Seed tuber microbiome is a predictor of next-season potato vigor

**DOI:** 10.1101/2024.04.18.590071

**Authors:** Yang Song, Elisa Atza, Juan J. Sanchez Gil, Doretta Akkermans, Ronnie de Jonge, Peter G.H. de Rooij, David Kakembo, Peter A.H.M. Bakker, Corné M.J. Pieterse, Neil V. Budko, Roeland L. Berendsen

## Abstract

Potato vigor, an important agronomic trait, is heavily influenced by the field of seed tuber production. Soil microbiota vary significantly between fields, impacting plant health and crop yield. Our study demonstrates that seed potato vigor can be predicted based on microbiota associated with seed tuber eyes, the dormant buds that grow out in the next season. By combining time-resolved drone-imaging of potato crop development with microbiome sequencing of seed tuber eyes from 6 varieties produced in 240 fields, we established correlations between microbiome fingerprints and potato vigor parameters. Employing Random Forest algorithms, we developed a predictive “Potato-Microbiome Informed” model, revealing variety-specific relationships between seed tuber microbiome composition and next season’s potato vigor in trial fields. The model accurately predicted vigor of seed tubers to which the model was naïve and pinpointed key microbial indicators of potato vigor. By connecting variety-specific microbiome fingerprints to crop performance in the field, we pave the way for microbiome-informed breeding strategies.

## Main

The microbial community inhabiting and surrounding plants, the plant microbiome, has long been recognized for its significant impact on plant growth and health ^1^. While specific microbes can cause disease, others play a vital role in supporting plant growth by effectively mobilizing scarce soil nutrients and protecting plants from pests and diseases ^2–4^. Plants try to shape the plant microbiome to their advantage ^5^ and can recruit protective microbes in response to pathogen or insect attack ^6^. As a result, plants can assemble plant protective microbiomes that survive in soils to protect subsequent plantings. In contrast, the buildup of pathogen inoculum in soils can also result in negative soil feedback and reduce growth of later plant generations ^7^. Plant microbiomes that affect plant performance can thus be inherited from one plant generation to the next, which is especially relevant for crops that are propagated via organs from the soil, such as potato tubers.

Potato is the world’s third most important crop for human consumption, valued not only as a food source but also as a significant source of starch for the pharmaceutical, textile and paper industry (FAO, 2017). Due to its high space and water use efficiency, potato yields five times more consumable weight per hectare than rice and wheat, making it crucial for global food security, as recognized by the UN-FAO ^8, 9^. Unlike most other major field crops, potatoes are primarily propagated vegetatively by transplanting seed tubers from one field to another in the next growing season.

Substantial variation in the growth potential of seed potatoes, also known as potato vigor, is observed when seed potato tubers of the same variety originate from different production fields. Potato vigor refers to the physiological potential for rapid and even outgrowth, good emergence, and proper development of new shoots sprouting from the tuber eyes ^10^. Eyes are the dormant buds or growth points present on the surface of a potato tuber from which new shoots sprout during the early stages of potato plant growth. Potato vigor is the highest at the end of a dormancy phase, after which it declines with the aging of the tubers during storage. In addition to age, vigor depends on the genetic background of a potato, as both the length of the dormancy period and the subsequent rate of aging differ between potato cultivars ^11^. Potato vigor is also strongly influenced by the growth history of the mother plants that produced the seed tubers ^12, 13^. When seed tubers of the same variety but from different production fields are planted together in the next season, abiotic and biotic differences in the field of production can lead to varied physiological and microbial imprints in the seed tuber, resulting in differential effects on potato vigor ^14, 15^. Crop yield and quality ultimately depend on potato vigor, and low vigor can significantly affect the livelihood of potato growers.

However, a reliable and rapid diagnostic tool capable of assessing differences in potato vigor does not currently exist ^11^.

Studies have demonstrated the significant impact of the potato tuber microbiome on plant health and productivity ^16–18^. While potato is susceptible to various plant pathogens that can be carried by the seed tuber ^19, 20^, the tuber can also host beneficial microbes that promote plant growth ^16, 17, 21^. Moreover, tubers produced in different fields harbor distinct microbiomes that can impact the microbiome of potato roots and even new tubers in a subsequent growing season ^14, 22, 23^. Here, we investigated whether the field-specific microbiomes carried by the tuber are associated with potato vigor, as exemplified by the extent of outgrowth of new shoots during early stages of plant growth in the new growing season. We employed microbiome-amplicon sequencing of a large set of field-collected tubers and extensive time-resolved drone-based imaging of potato crop development in the next season to build a model that can reliably predict vigor of seed potatoes. Moreover, our model identifies microbial sequences that are potato-variety-specific predictors of potato vigor.

## Results

### Field of production affects seed tuber vigor

Variability in potato vigor caused by physiological and microbial imprinting of the seed tuber in its field of production is a major issue in the potato production system. To establish the extent of variation in vigor of the emerging potato plant in the next growing season, we collected seedlots (batches of seed potato tubers that each originate from a single production field) of 6 different potato varieties, each variety harvested from 30 seed tuber production fields across the Netherlands in both the autumn of 2018 (year 1) and 2019 (year 2; Fig. 1A, Fig. S1). In two consecutive years, seed tubers from 180 seedlots were stored over winter and planted in each of 3 trial fields in the next spring. In both years, the 3 trial fields were located on farms near Montfrin (M) in France and near Kollumerwaard (K) and Veenklooster (V) in the Netherlands (Fig. S1C). In each of the trial fields, the seed tubers were planted in a randomized block design with 4 replicate blocks of 24 tubers evenly distributed over 4 ridges (Fig. 1B, C, Fig. S1). We monitored the growth and development of the plants that emerged from these seed tubers using aerial images of the complete field with a drone-mounted camera from 30 days after planting (DAP) until 50 DAP, when leaf canopies of neighboring plants began to overlap and there were no more empty ridges detected (Table S1). As a measure for potato vigor, we corrected for spatial heterogeneity in the field (Fig. S2) and quantified the canopy surface area (CSA) of each replicate block (Fig. 1C).

**Fig. 1.**
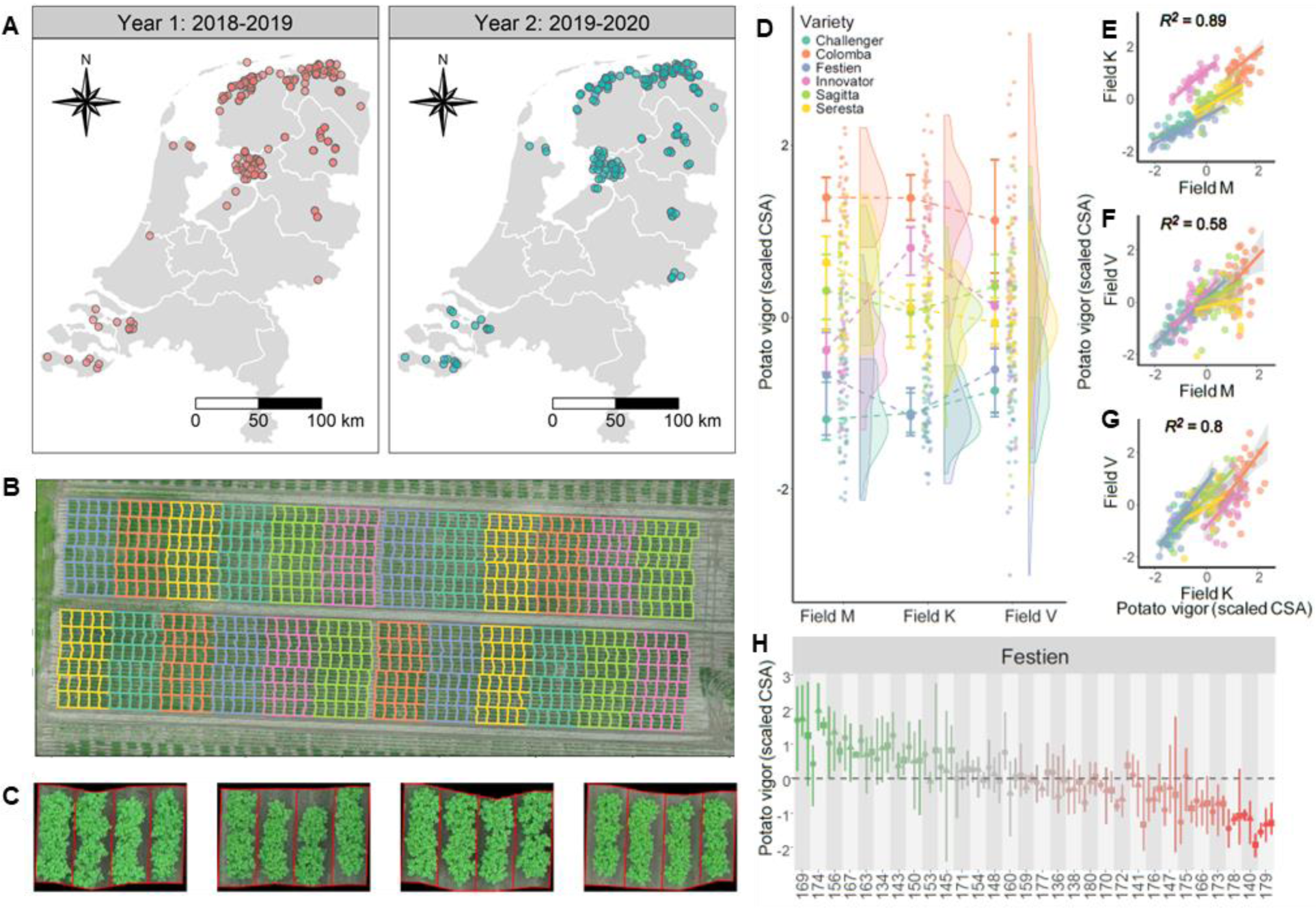
Variation in canopy development of potato plants from 6 varieties and 180 production fields. **A** Locations of the 180 fields per year where the seedlots of 6 potato varieties (30 fields per variety) were collected in the Netherlands. **B** Drone image showing the complete randomized block design in trial field M in year 2. All 30 seedlots of the varieties Challenger (green), Colomba (salmon), Festien (blue), Innovator (mauve), Sagitta (green) and Seresta (yellow) were planted in 4 replicated blocks. Within a block, tubers of each seedlot were planted in a plot of 4 by 6 tubers. **C** Exemplary images of 4 replicate plots of the same seedlot in trial field M. **D** shows the distribution of scaled CSA among trial fields on 52 (field M), 48 (field K) or 50 (field V) days after planting in year 1. **E-G** Spearman correlations of scaled CSA across fields. *R*^2^ values result from fitting a linear mixed model to scaled CSA using plant varieties as random effects. The *ρ* and significance of Spearman correlations of scaled CSA across fields of all varieties or per variety are depicted in Table S2. **H** Effect of each seedlot on the growth of Festien plants in each of the three trial fields in year 1. CSA is scaled to the variety average in each trial field. Symbols and error bars show average, minimum and maximum CSA values of 4 replicate plots of each seedlot per trial field, after correcting raw CSA values by spatial heterogeneity.

In both years and all 3 trials fields, we observed that the CSA of the 6 potato varieties developed at substantially different variety-specific growth rates (Fig. 1D; Fig. S3), but also that there was significant variation in the performance of the distinct seedlots within each variety (Fig. 1E-G; Fig. S4). We calculated Spearman correlations to assess the relationship between CSA of different seedlots in each trial field and found a strong overall correlation (*P* < 0.001) for all pairwise comparisons of CSA across trial fields in both experimental years (Fig. 1E-G, Table S2), showing that potato vigor of a seedlot in one trial field is correlated to the vigor of the same seedlot in another trial field. For the 6 varieties independently, 31 out of 36 pairwise comparisons of seedlot CSA were significantly correlated (Table S2). Three out of the 5 non-significant correlations, in field V during the second year, were likely influenced by an herbicide treatment, which especially affected the early-emerging varieties Colomba and Innovator. The vast majority of the correlations thus were significant, and this suggests that the rate of canopy development is an intrinsic property of the seedlot, and that potato vigor affects plant outgrowth in the next season. For variety Festien (Fig. 1H, Fig. S4), as well as other varieties (Fig. S5), this seedlot-dependent variation in potato vigor is consistent in all 3 trial fields as is visible when CSAs was normalised for the average of the field (scaled CSA). Seedlots consistently perform above (green) or below (red) the variety’s average vigor. Together these data suggest that rate of canopy development reflects the potato vigor of a tuber seedlot at time of planting in the trial field and that at least part of potato vigor is imprinted during growth of the mother tuber in the production field.

### Edaphic factors in the field of production and plant genotype affect seed tuber microbiome

To investigate if the variation in potato vigor between seedlots is associated with differences in the composition of the tuber microbiome, tuber eyes and heel ends were sampled from tubers in the months after harvesting. Whereas 180 seedlots were harvested in each year for the potato vigor assessments (Fig. 1), we selected a subset of 60 seedlots from year 1 for seed potato eyes and heel ends microbiome analysis (Table S3) and used 4 replicate samples of 50 tubers per seedlot (240 samples in total for year 1). Preliminary analysis showed that replicate samples comprised very similar microbiomes, so for microbiome sequencing in year 2 we used only 2 replicate samples per seedlot, but we analysed the microbiomes of tuber eyes and heel ends from tubers of all 180 seedlots (360 samples in total for year 2; Fig. S1). From those 600 samples of tuber eyes and of tuber heel ends from 240 distinct seedlots across 2 experimental years, we subsequently amplified and sequenced 16S rRNA and ITS regions of respectively the bacterial and fungal DNA (Fig. S1). After clustering sequencing reads into amplicon sequence variants (ASVs) and filtering out ASVs occurring in less than 2 samples, we identified a total of 17874 bacterial ASVs and 1755 fungal ASVs in the eye compartments, and 20119 bacterial ASVs and 1917 fungal ASVs in the heel end compartments of tubers.

To identify microbiome fingerprint signatures in the microbial communities of the seedlots across soil type, potato variety, and harvesting year, we computed the Bray-Curtis dissimilarity for all pair-wise sample combinations and performed a principal component analysis (PCoA). A permutational multivariate analysis of variance (PERMANOVA) on the tuber eye data revealed significant (*P* = 0.001) effects of soil type, variety, and year on the composition of bacterial (Fig. 2A-C) and fungal (Fig. 2D-F) tuber eye microbiomes. Of these factors, soil type and variety had the strongest impact on the bacterial community, accounting for respectively 11% and 12% of the observed variation, while 2.0% of the variation in microbiome composition was related to the year of harvest. This suggests that the tuber eye microbiome is shaped by a combination of edaphic factors and plant genotype. Moreover, when examining the microbiomes of tuber eyes from replicates from each seedlot, we found that tuber eyes from the same seedlot have similar microbiomes and are significantly different (*P* < 0.01) from those of other seedlots (Fig. S6, Fig. S7). Similar results were observed for the heel end compartment (Fig. S8). This again underlines that the field of production shapes the seed tuber microbiome ^14^.

**Fig. 2.**
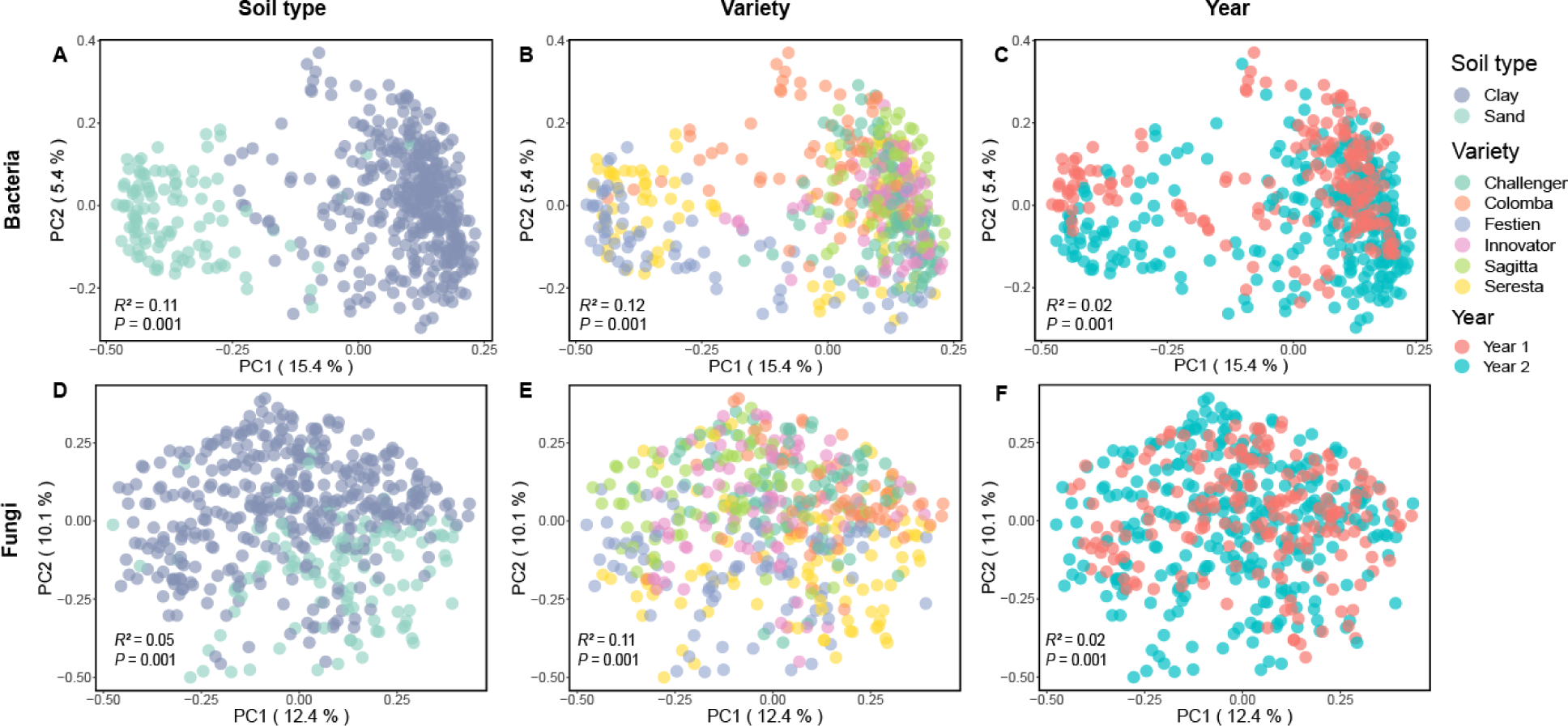
Beta diversity analysis of seed tuber microbiomes in the tuber eye compartment. PCoA ordination plot based on Bray-Curtis dissimilarities of bacterial (**A-C)** and fungal **(D-F)** microbiomes are colored by soil type (**A,D**), variety (**B,E**) and year (**C,F**) as indicated in the legend. Each data point represents a single replicate of a seedlot. Four replicate samples were analyzed for each of 60 seedlots in year 1 and two replicate samples for each of 180 seedlots in year 2. *P* and *R*^2^ in each PCoA are the result of PERMANOVA on soil type (**A,D**), variety (**B,E**) and year (**C,F**) as respective factors.

### Microbiome-based predictions of seed tuber vigor

Having established that for each variety seedlots clearly vary in both vigor and microbiome composition, we investigated whether we could construct a model that is able to predict seedlot potato vigor using microbiome fingerprint data from the eyes of the seed tubers. Lately, machine-learning techniques have been successfully applied to microbiome data ^24–26^ and we constructed a microbiome data-based Random Forest model ^27^ to predict potato vigor. To first determine the optimal taxonomic rank for the prediction, we assessed model performance using distinct taxonomic levels - i.e., ASV, species, genus, family, order, and class - as features for the model. Moreover, we used hierarchical feature engineering (HFE) ^28^ to generate an additional feature level. HFE is a method used for feature engineering in machine learning that incorporates hierarchical information into the feature representation. It leverages the hierarchical structure of microbiome data and selects features of different taxonomic rank to optimize the prediction.

We trained the model on the bacteria data of year 2 and potato vigor data (CSA) in the trial field M in the same year (training set). To avoid predicting differences in CSA resulting solely from genetic differences between potato varieties, we used scaled CSA values per trial field and variety to train our RF model to minimize the error in fitting out-of-bag (OOB) samples. Based on the OOB coefficient of determination (*R*^2^), HFE initially stood out as the preferred option to predict plant vigor, followed by ASV (*R*^2^=0.34 and 0.22, respectively; Table 1).

**Table 1.**
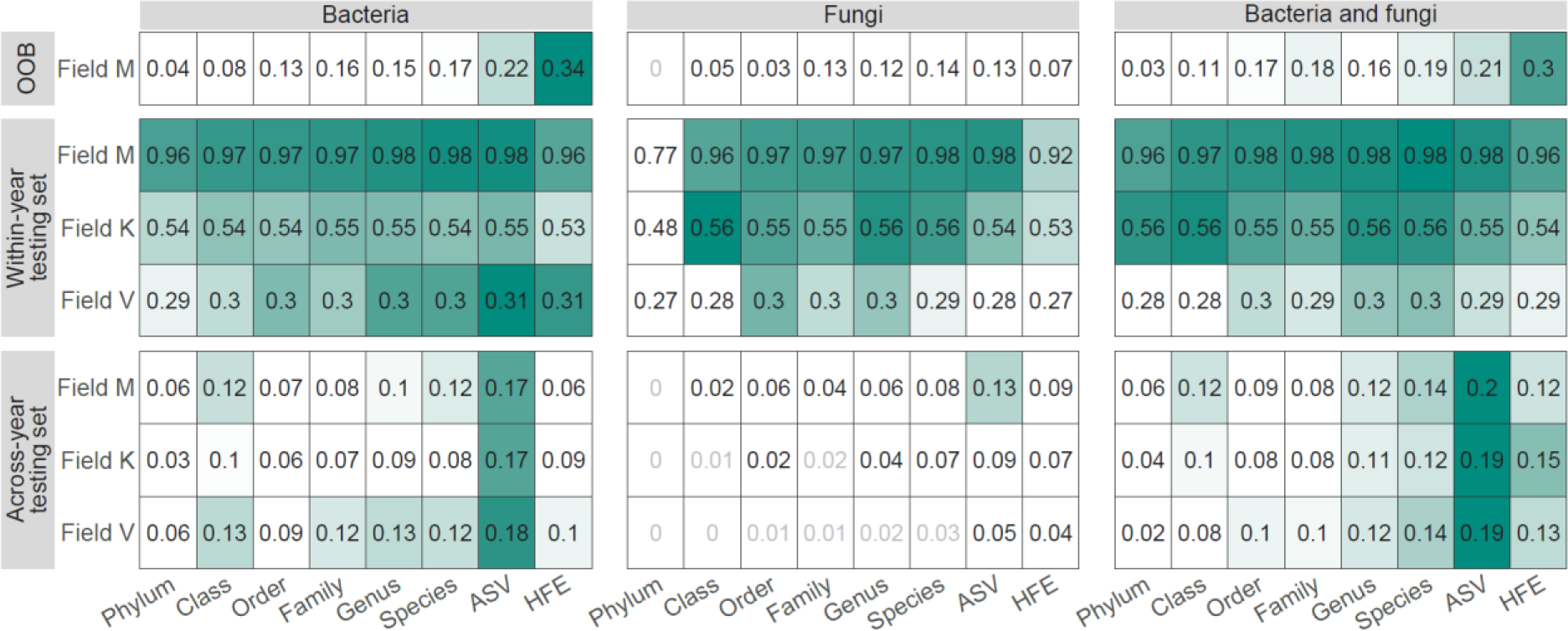
Performance of RF models trained with tuber eye microbiome data at distinct taxonomic ranks and HFE features. Random Forest models trained on tuber microbiome data and Field M potato vigor in year 2 were tested on vigor data from field M (training set), field K and V in year 2 (within-year testing set) and then tested on data from field M, K and V in year 1 (across year testing set). Numbers represent the *R*^2^ of a model, where a higher value indicates superior performance. Numbers in black represent significant correlations where *P* < 0.05, gray represent unsignificant corrections. OOB indicates the out-of-bag performance of the models in the training set.

We then tested the model (trained on observations in Field M) by correlating the predicted CSA to the CSA observed in the two other trial fields of the same year (within-year testing set). We found that the potato vigor-related CSA values predicted by the model based on field M were significantly correlated to those observed in the other two trial fields of year 2, regardless of taxonomic rank of the bacterial data (within-year testing set; Table 1). For within-year predictions, training the model on HFE resulted in similar *R*^2^ values as when the model was trained on ASV features. Within-year predictions, however, are all based on microbiome data from the same year. We then tested the Random Forest model further with the tuber microbiome data from year 1 (across-year testing set), to which our model was completely naïve. Although the prediction performed better for fields within the same experimental year, even the prediction based on the seedlot microbiomes from this preceding year that were not used to train the Random Forest model were significantly correlated to the observed vigor of the seedlots. Here, the HFE-based Random Forest model (maximum *R*^2^ = 0.1; Table 1; Bacteria) was outperformed by the Random Forest model trained at the ASV level (maximum *R*^2^=0.18).

Subsequently, we compared the performance of Random Forest models that were built on either bacterial data alone (Table 1; Bacteria), fungal data alone (Table 1; Fungi) or data that combined bacterial and fungal microbiomes (Table 1; Bacteria and fungi). Training on fungal data alone resulted in reasonably similar performance of the model with within year data but in a lower model performance in the across-year testing set compared to the model based on only bacteria at any phylogenetic level (Table 1B). Combining bacterial and fungal ASV tables slightly improved the predictions of the bacteria-only Random Forest model for both the within-year and across-year testing sets (Table 1C).

Collectively, our results demonstrate that our “Potato-Microbiome Informed” (PMI) predictive model based on microbiome fingerprints of seed tubers can predict potato vigor in the next growing season. The PMI model performs most accurately across-year when it is based on the highest taxonomic resolution at the ASV level and improves when based on data from both bacterial and fungal amplicons. We also successfully trained Random Forest models on heel end microbiome data (Table S4). The performance of the models based on the heel end microbiome data were slightly less accurate than the models based on tuber eye microbiome data, and we focused on the latter (henceforth: tuber microbiome) for the remainder of this study.

### Practical application of the microbiome-informed prediction model for potato vigor

To potato producers, seed tubers represent a large yearly investment, and hence, their vigor is of considerable importance. The potato vigor CSA values predicted by the PMI model correlate significantly with the observed CSA values both for the within-year testing set (Fig. 3A; Spearman correlation) as well as for the across-year testing set (Fig 3B).

**Fig. 3.**
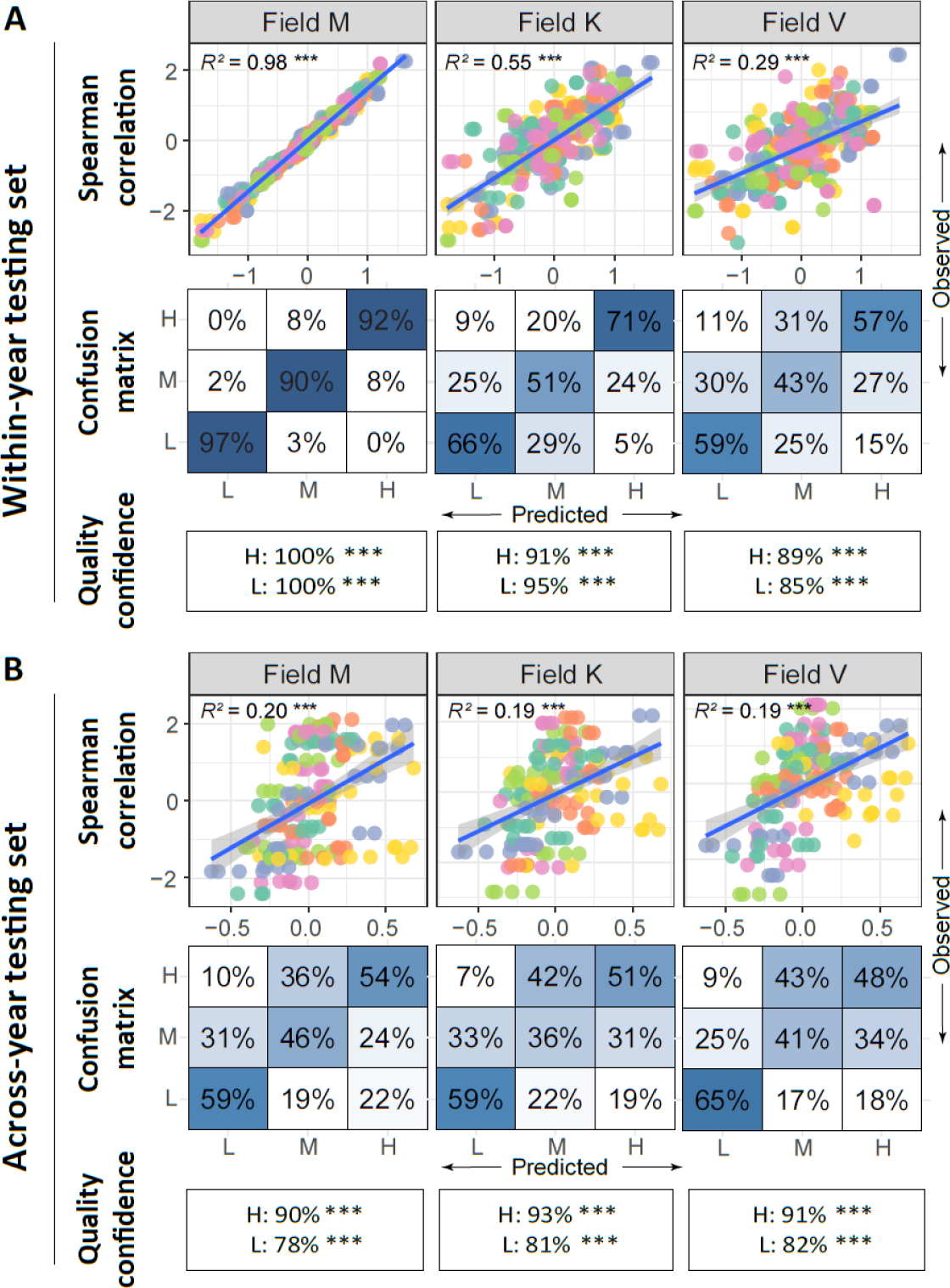
Model performance on within-year and across-year test sets over all varieties. Model performance of all varieties and per variety on **A** within-year testing sets and **B** across-year testing sets. Scatter plots illustrate the Spearman correlation between the predicted potato vigor (x-axis) and observed potato vigor (y-axis). Predicted values are generated by the PMI model trained on microbiome data from year 2 and CSA from field M of year 2. Predicted CSA is either based on the same microbiome data (within-year) or on microbiome data from year 1 to which the model was naïve (across-year). Predicted and observed CSA are scaled to the variety average in each trial field. Confusion matrices showing the precision of the model in identifying classes of high (H), medium (M) and low (L) CSA. Quality confidence of within-year and across-year predictions, i.e. the chance of respectively the H or L class not being misclassified as the opposite extreme class. Asterisks indicate pseudo-*P* (\**P* < 0.05; \*\**P* < 0.01; \*\*\**P* < 0.001) of quality confidence being higher than random guess by simulating 1000 random classifications.

To intuitively evaluate the performance of the PMI prediction model, we divided the seedlots into three classes based on their predicted and observed vigor (CSA value): high vigor (H; 1/3 of the seedlots with highest vigor), low vigor (L; 1/3 of the seedlots with lowest vigor) and medium vigor (M; rest 1/3 of the seedlots). We then calculated the precision value of the model (Fig. 3; Confusion matrix), which represents the percentage of correctly predicted samples within each respective class. Whereas the model predicts L or H classes relatively accurately, the model appears to have more difficulty distinguishing M from L or H, especially in the across-year testing set (Fig. 3B; Confusion matrix). Since users will be most-interested in knowing the chance that a seedlot that is predicted to have high vigor is not in reality a seedlot with observed low vigor, or the other way around, we defined the estimated chance of a seedlot to be predicted in one of the two extreme classes H or L, but is misclassified in the opposite (L or H) as the “quality confidence” of the prediction (Fig. 3; Quality confidence). In the within-year testing set, this yielded a high quality confidence score of 85 – 100% of the seedlots that were predicted to be H or L in reality not being the opposite (L or H). Interestingly, even in the across-year testing set, only a small percentage (5 - 22%) of seedlots were misclassified and predicted in the opposite class. For instance, of the seedlots that were predicted to have high vigor, 90-93% had a high or medium observed vigor and only 7-11% were misclassified as low vigor. Conversely, seedlots that were predicted to have low vigor, 78-82% had indeed an observed low or medium vigor and only 18-22% was misclassified as high vigor, which is significantly lower than random (*P* < 0.001, Fig. 3B). Even though the precision values of each class across years (Fig. 3B; Confusion matrix) are generally lower than the precision values within year (Fig. 3A; Confusion matrix), the quality confidences indicate that the model still demonstrated favorable performance in predicting the H or L class. This shows that the prediction model is fairly accurate in predicting potato vigor extremes, and that the model has real-world applicational value in assessing potato vigor. accurate in predicting potato vigor extremes, and that the model has real-world applicational value in assessing potato vigor.

### Performance of the model differs across potato varieties

The model described above was designed as a variety-generic model trained on data from all 6 potato varieties used in this study. When looking into each individual variety, within the same year, the potato vigor CSA values predicted by the model correlate significantly with the observed CSA values for each of the 6 varieties. Although the quality confidence for each of the varieties varies, it is generally high (82-100%) within year (Fig. 4A).

**Fig. 4.**
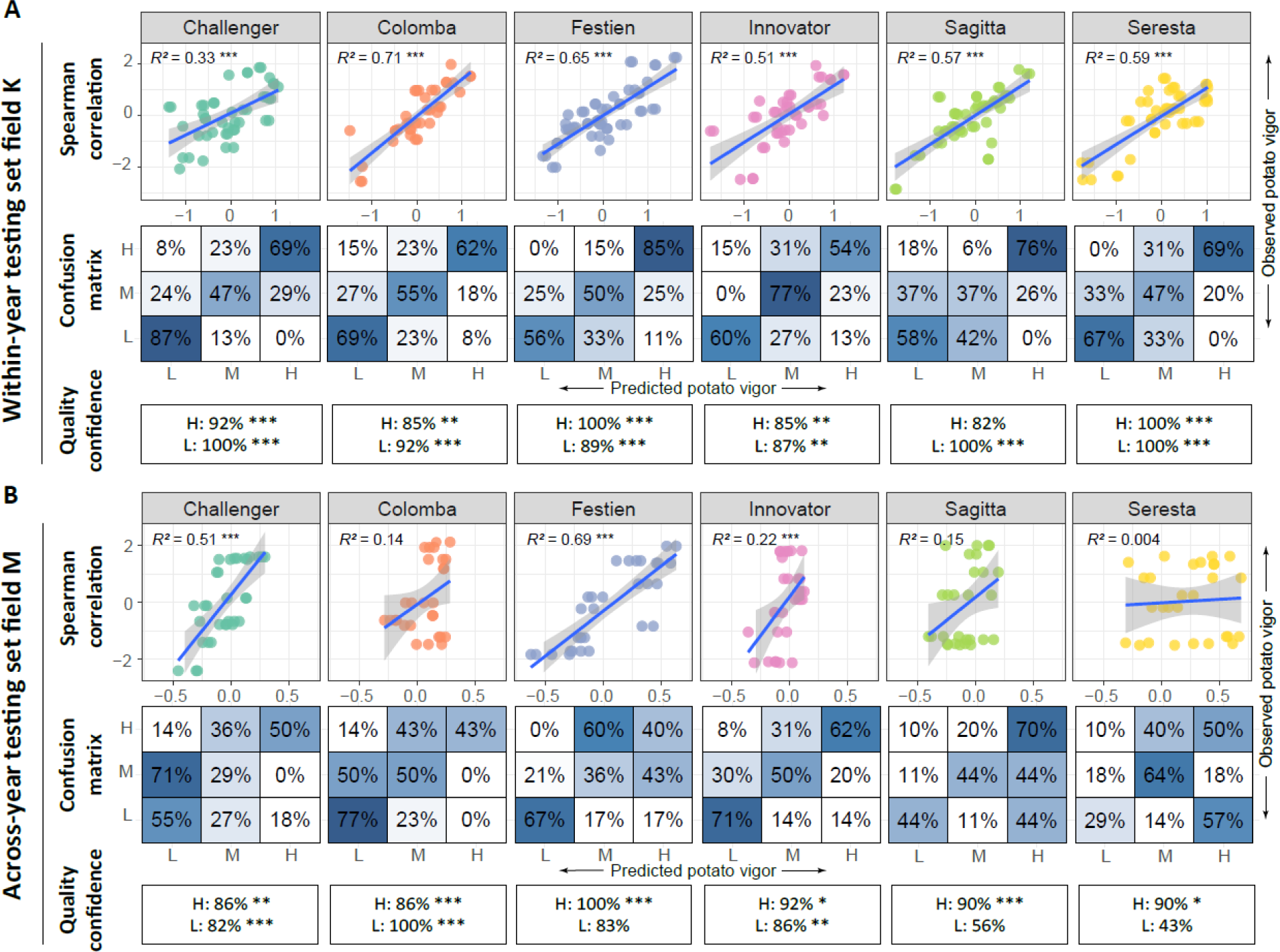
Model performance per variety on within-year and across-year test sets. **A** Model performance per variety on within-year test sets. **B** Model performance per variety on across-year test sets. Scatter plots illustrate the Spearman correlation between the predicted potato vigor (x-axis) and observed potato vigor (y-axis). Predicted values are generated by a random forest model trained on microbiome data from year 2 and CSA from field M of year 2. Predicted CSA is either based on the same microbiome data (within-year) or on microbiome data from year 1 to which the model was naïve (across-year). CSA is observed in Field K in year 2 (**A**; within-year), or in Field M, year 1 (**B** across-year). Predicted and observed CSA are scaled to the variety average in each trial field. Confusion matrices showing the precision of the model in identifying classes of high, medium and low CSA. Quality confidence of within-year and across-year predictions, i.e. the chance of respectively the H or L class not being misclassified as the opposite extreme class. Asterisks indicate pseudo-*P* (\**P* < 0.05; \*\**P* < 0.01; \*\*\**P* < 0.001) of quality confidence being higher than random guess by simulating 1000 random classifications.

The microbiome-based prediction of potato vigor across years, however, revealed bigger differences in model performance across different potato varieties. Spearman correlation analysis indicated a significant positive correlation between predicted and observed vigor for the varieties Challenger, Festien and Innovator, which indicated that the model exhibited stronger predictive performance for these varieties (Fig. 4B, Fig. S9). The across-year prediction was less informative for varieties Colomba, Sagitta and Seresta. This shows that the PMI model does not work equally well for all varieties. Nonetheless, even though the correlations between observed and predicted potato vigor values are not significant for Colomba, Sagitta and Seresta, the quality confidence parameter still provides relatively high scores (86-90%) for the high-vigor seedlots classified as H (Fig. 4B).

### Key microbial predictors of potato vigor and their variety-specificity

The predictions described above are based on the occurrence and abundance of 19629 bacterial and fungal ASVs detected on the eye compartment of potato tubers across the Netherlands. In order to identify the microbial taxa representing the key predictors of the PMI potato vigor prediction model, we ranked ASVs by their contribution to the prediction and selected the top ASVs that together accounted for either 1% (20 bacterial and 2 fungal ASVs) or 5% (96 bacterial and 14 fungal ASVs) model’s accuracy (Fig. 5A). Among the key predictors, the majority of the ASVs corresponded to bacterial taxa belonging to Pseudomonadota, Actinomycetota or Bacteroidota (Fig. 5B). The abundance of these ASVs exhibited both positive and negative correlations with seed tuber vigor (Fig. 5C-E; Fig. S10; Fig. S11). The 3 ASVs that contribute most to the prediction of the PMI model are *Streptomyces* ASV 6c8e8, *Acinetobacter* ASV bcae8 and *Cellvibrio* ASV c205d. Whereas the abundance of *Streptomyces* ASV 6c8e8 on the tuber is positively correlated with tuber vigor, higher abundances of *Acinetobacter* ASV bcae8 and *Cellvibrio* ASV c205d are negatively correlated with potato vigor (Fig. 5C-D).

**Fig. 5.**
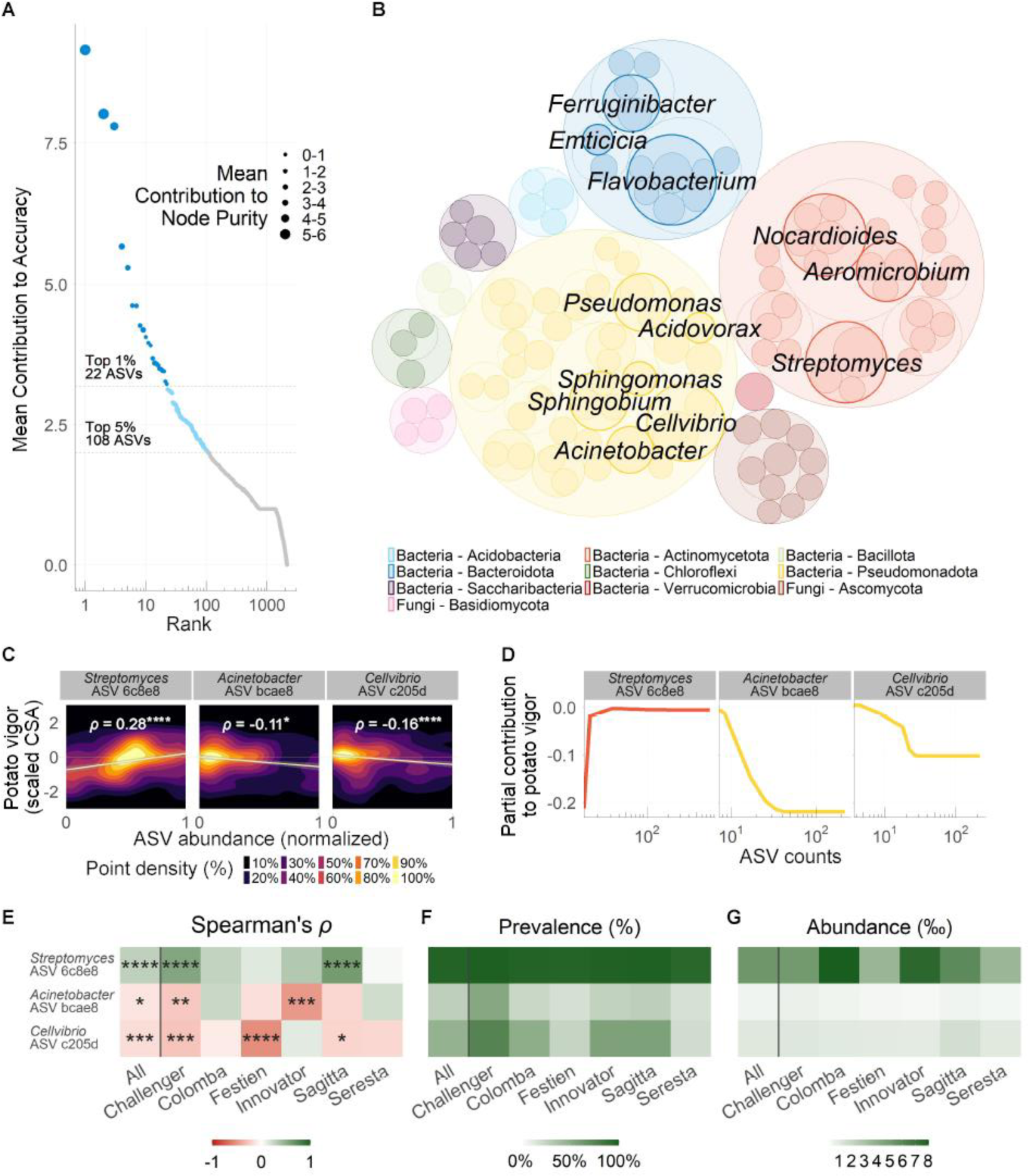
The most predictive microbes selected by the RF prediction model. **A** Mean contribution to accuracy of the RF model of each of the ASVs, ranked from highest to lowest contribution. The top contributing ASVs are highlighted representing respectively 1% (dark blue color, 22 ASVs) and 5% (light blue color, 86 additional ASVs) contribution to the accuracy of our model. **B** Bubble plot depicting the phylogenetic composition of the top 5% most-predictive microbes, grouped, and colored according to their phylum. The top 1% most-predictive ASVs are highlighted with their genus classification when present. Bubble size indicates the mean contribution to the accuracy. **C** Spearman correlation between the relative abundance of each top 3 contributing ASV and potato vigor. ASV abundance is rescaled between 0 and 1 with respect to their minimum and maximum to show one single scale across ASVs. Color represents the density of the sample points. The clearest colors indicate areas that accumulate most of the samples, and dark colors the areas where no data or few sample points are found. The line represents robust regression to outliers and Spearman’s *ρ* is shown, whereas asterisks indicate significance level (\**P*<0.05; \*\*\*\**P*<0.0001). **D** Partial contribution plots for the top 3 ASVs most predictive to plant vigor according to the RF model. **E** Heatmaps showing Spearman correlations of each of the top 1% contributing ASVs to plant vigor, their prevalence, and median abundance across samples. The first column in every heatmap shows the computed value including all the data regardless of plant variety, and the rest of columns display those values calculated for individual potato varieties.

One of the main advantages of applying a Random Forest algorithm is that the reported important ASVs do not necessarily need to correlate linearly with potato vigor CSA values. Instead, the PMI model can capture more complex relationships that can be examined with partial contributions. The partial contribution of an ASV represents how the predictions of potato vigor change according to the counts of that individual ASV. To explore the relationship of each of the top 1% ASVs with potato vigor, we computed the partial contributions of each ASV to the model predictions (Fig. S10B).

*Streptomyces* ASV 6c8e8 was the best predictor of vigor. The vigor of the tubers in which this particular ASV was not detected was below average, but, on the tubers that did have this ASV, increased abundance of ASV 6c8e8 was associated with improved potato vigor. In addition, while this *Streptomyces* ASV is both prevalent and fairly abundant in most microbiome samples (Fig. 5E), Fig. 5D shows that above a threshold level of approximately 30 counts per sample potato vigor is not further improved. This suggests that especially the absence of *Streptomyces* ASV 6c8e8 on the seed tuber is a crucial indicator of low potato vigor. In contrast, the presence of *Acinetobacter* ASV bcae8 and *Cellvibrio* ASV c205d are indicators of relatively low potato vigor. Moreover, reduced vigor of the tubers correlated with increased abundance of these ASVs. Remarkably, the negative correlation between abundance of these ASVs and vigor of the tubers appears to reach a plateau after which further increase in abundance of these microbes has no additional negative effect on potato vigor (Fig. 5D).

Whereas the key predictive microbes were identified by training the PMI model on data of all varieties, the correlation between abundance and potato vigor was not apparent on all of the individual potato varieties (Fig. 5E; Fig. S11). Such variety-specific relationships suggest that seed potato vigor of distinct varieties is linked to the presence and abundance of different key microbes. Together these results show that it is possible to identify key microbes that are predictive for the vigor of seed potato tubers and that these microbes can thus be developed as potential biomarkers for potato vigor.

## Discussion

The vigor of potato seed tubers, as reflected by the rate of canopy development of the emerging shoots, varies ^14, 29^. Seed tubers harbor a microbiome that is largely assembled in the field of seed tuber production and is carried to the field in which the tuber is planted ^14^. Microbes on a seed tuber likely partly reflect the physiological state of that tuber, but tuber-borne microbes can also affect the growth of the plant that emerges from a tuber ^30, 31^. Here we demonstrate that potato tuber vigor can be predicted by tuber microbiome fingerprint data.

### Vigor is imprinted in the field of production

After seed tubers are planted in a field, shoots grow out of the tuber’s eyes, gradually developing into a canopy. Our data demonstrate that distinct potato varieties exhibit varying rates of canopy development, highlighting the evident relationship between potato genetics and growth.

Additionally, climatic conditions and soil characteristics in the planting field also influence the rate at which a potato variety develops, resulting in varying average canopy surface areas (CSAs) for each variety in different trial fields. Nonetheless, our data show that even seed tubers from the same variety and grown in the same trial field develop canopies of different sizes, depending on the production field from which the mother tuber originated. Furthermore, we found that the CSA produced by a seedlot is correlated across trial fields, and that seedlots outperforming or underperforming the average of its potato variety, consistently do so across different trial fields. These seed tubers thus exhibit distinct potential for developing CSA, and we considered CSA to be a reflection of the tuber’s vigor at the time of planting. Since all seedlots in our study were stored under the same conditions in one location, our data underscore that potato vigor is, in part, imprinted in the seed potato tubers by local conditions in their production field.

### Each field produces a unique seed tuber microbiome

We analyzed the seed tuber microbiome composition of 240 seedlots across the Netherlands. Consistent with previous studies ^32, 33^, we found that soil type, year of harvest, and potato variety were significant factors affecting the microbial signatures found on the seed tubers. It is well-established that abiotic factors, including both edaphic and climatic conditions, influence the soil microbiome ^34^. Additionally, each field has a distinct pre-crop history, during which each crop species or even variety selectively assembles a distinct microbiome ^35^. Agricultural practices also play a role in shaping the microbiome composition of soil ^36^. Plants can selectively assemble a microbiome from the available microbes in the soil. Consequently, we found that seed tubers harbor a distinct microbial fingerprint, depending on the soil in which they were produced.

### Microbiome-informed prediction of crop vigor

We were able to link the composition of seed tuber microbiome to plant performance in the subsequent growing season. Utilizing amplicon sequence data of tuber microbiomes from one year, we generated a Random Forest-based PMI potato vigor prediction model that successfully predicted potato tuber vigor in a different year. This demonstrates that the microbial signature of the seed tuber can predict the outgrowth of emergent plants in the next season. Hence, our PMI model holds direct applicational value, serving as a diagnostic test to assess tuber seedlot quality. Additionally, the PMI model shows that machine learning has the potential to generate models for vigor assessment of seeds or seedlings of other plant species. In this regard, initial steps have been taken to identify soil microbiome indicators indicative of maize growth responses to inoculation of arbuscular mycorrhizal fungi ^37^. Also, the occurrence of Fusarium disease could be predicted based on soil microbiome composition ^26^.

### Key microbial predictors: cause or consequence of vigor

The PMI model not only predicts the vigor of seed tubers but also ranks the features contributing to the prediction. The key features identified by the model could be considered as biomarkers for vigor.

At this point, it is not possible to distinguish whether these ASVs are merely a reflection of the physiological state of the tubers or the cause of improved or deteriorated vigor. Nonetheless, there are some indications in the literature that argue at least some of the key microbes affect the vigor of the tubers. The presence of *Acinetobacter* ASV bcae8 is linked to decreased vigor of tubers by our model, which is consistent with observations that *Acinetobacter* sp. strongly reduces the growth of potato seedlings ^38^. Moreover, the feature contributing most to the prediction of our model is *Streptomyces* ASVs 6c8e8, the presence of which appears to be a prerequisite for good vigor. In that respect, it is remarkable that the natural buildup of high densities of pathogen-antagonistic *Streptomyces* spp. in soils is thought to play a key role in natural suppression of potato disease ^39–42^. Therefore, our model is also a potential treasure trove for the discovery of unrecognized deleterious or beneficial microbes specific to potatoes. Our current efforts are focused on isolating and identifying the microbes that represent the keystone ASVs from potatoes and validating their effects on potato vigor.

### A working model and avenues for improvement

When the PMI model was applied to predict potato vigor based on microbiome data of seed tubers to which the model was naïve, a maximum variation of 20% was explained by the model. This implies that the tuber microbiome explains only part of the seed tuber vigor, and the model could be improved by incorporating additional variables such as soil properties, field planting history, and the metabolomic properties of the planting material. The prediction could also be improved by using different machine learning algorithms such as gradient boosting ^43^ and neural networks^44^. In this study, Random Forest ^27^ was chosen due to its inherent advantages in handling high-dimensional data characteristic of microbiomes. Random Forest has been widely used in microbiome host trait prediction in clinical studies and showed superior performance to other models ^33, 45^. Additionally, in environmental studies, Random Forest has demonstrated the best performance in predicting the disease occurrences in soils by sequencing soil microbiomes ^26^. Moreover, we chose a Random Forest regression approach to enable the derivation of a relative vigor score, and to empower users to accurately pinpoint seedlots with high or low vigor. Thus, the vigor categories are more adaptable and fitting for practical applications. Finally, we used HFE in this study ^28^ for feature selection and Random Forest default hyperparameters to train the models. Other methods for feature selection (e.g. Fizzy ^46^ and MetAML^45^) and hyperparameter tuning are promising avenues to improve the performance of the PMI model ^47^.

### Towards microbiome-optimized potato

Finally, we found that the vigor of distinct potato varieties is linked to specific subsets of the identified predicting microbes. This observation aligns cohesively with existing studies that highlight divergent sensitivities among plant varieties to both pathogens ^48^ and beneficial microorganisms ^49^. Our data and similar approaches linking microbiome composition to plant phenotypes could be used to unearth interactions of beneficial and deleterious microbes with specific genotypes, genetic regions and ultimately even genes of potato and other crops. This would be the first step in the process of breeding or engineering elite crop varieties that produce more but require less, as they consistently assemble better-functioning microbiomes. This transformative insight not only advances our understanding of microbial contributions to plant performance but also lays the groundwork for a microbiome-based breeding strategy aimed at enhancing the quality of planting material.

### Online Material and Methods

#### Potato varieties and seedlots

In total, 6 potato varieties form the Royal HZPC Group and Averis Seeds B.V. were used in this study, namely variety Challenger (Variety A), Colomba (Variety B), Festien (Variety C), Innovator (Variety D), Sagitta (Variety E) and Seresta (Variety F). In the autumn of 2018 (year 1) and 2019 (year 2), we collected batches of seed tubers (seedlots) of 6 potato varieties from 30 different fields per variety (180 fields in total) in the Netherlands (Fig. 1A, Fig. S1). The 360 seedlots (180 in year 1 and 180 in year 2) were shipped to a central location where they were subsequently stored in the dark at 4 °C.

### Field trials setup and vigor measurement

Tuber seedlots from 180 fields per year were stored over winter after which they were planted in each of 3 trial fields in the following Spring. In both years, the trial fields were located near Montfrin (M, 54.4980 N, 5.1090 E) in France and near Kollumerwaard (K, 70.4325 N, 6.9825 E) and Veenklooster (V, 70.3935 N, 6.7080 E) in the north of the Netherlands, respectively. In each of the trial fields, the seed tubers were planted in randomized block design with 4 replicate blocks of 24 tubers evenly distributed over 4 ridges (Fig. 1B and C, Fig. S1). We monitored the growth and development of the plants that emerged from these seed tubers using aerial images of the complete field with a drone-mounted camera. These images were acquired starting from approximately 30 days after planting (DAP) of the seed tubers and continued until around 50 DAP when no more empty ridges were detected (Table S1).

### RGB image post-processing and canopy measurement

Due to the relatively large scale of the trials and the chosen planting technique, the trial fields did not exhibit the usual regular structure with easily identifiable rows and columns of plots. Therefore, an in-house standardized procedure was developed with minimal manual interaction to detect and identify plots in the trial fields’ images. The plot-detection algorithm is defined by 6 main steps. Step 1: From the provided row-column plot scheme, the expected number N of plots along the ridges of the trial fields is identified. Step 2: The field image where the canopy size allows to detect the gaps between the plots along the ridges is selected from the available images of the trial field. Such an image is usually found towards the end of the canopy growth season, where canopies inside a plot are touching, but have not yet grown as to bridge the gaps to neighbouring plots. Step 3: The beginning and the end of the trial field along each ridge is interactively determined in the selected image. Step 4: The expected number of N − 1 inter-plot gaps are automatically detected in the images along each ridge. Step 5: The detected plot polygons are displayed and inspected for possible remaining inaccuracies and distortions and the wrongly identified plot boundary points are corrected interactively. Step 6: For each plot a set of image coordinates of the plot polygonal boundary is saved.

To measure the canopy area within the polygonal plot boundary, the image pixels are segmented into two disjoint sets: pixels of the canopy and pixels of the surrounding soil. Then, the canopy pixels are counted, and the result converted to the cm^2^ units. The present dataset featured a variety of illumination and moisture conditions, both of which affect the color of pixels. Also, leaf canopy colors have systematic differences between varieties, ranging from light green to almost purple. Therefore, every orthophoto had to be processed individually, resulting in different segmentation filters with date- and field-specific parameters (see ref for details). In all cases, the quality of segmentation has been confirmed by visual inspection of randomly selected plots of each genotype.

After segmentation, the mean canopy area *S*px (in pixels) over each ridge of each plot was determined by summing all white pixels within the geometrical boundaries of the ridge and dividing by 6 – the number of plants in each ridge. A canopy area in pixels is converted to its area in cm^2^, using the distance *d*cm between the ridges in the field as was determined by the planting device and recorded as *d*cm = 75 cm in the Veenklooster (V) and Kollumerwaard-SPNA (S) fields, and *d*cm = 74 cm in the Montfrin (M) field. To find the pixel-to-cm conversion factor, we compute the average pixel distance *d*px between the adjacent ridges in the field for a specific date (see protocol Figure 2). Then, the area *S*1 of a single pixel in cm^2^ is given by:

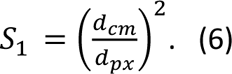

Thus, the canopy area *S* in cm^2^ is obtained from the canopy area *S*px in pixels as:

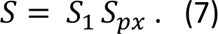

The ridge-mean plot canopies obtained after the transformation, plot localization, and segmentation procedures described above constitute the raw data and cannot be used to estimate the mean batch canopy, since the test fields are usually spatially inhomogeneous, which may systematically increase or decrease the canopy size in certain areas of the field. We used the state of the art spatial effect removal method implemented in the R-package SpATS ^50^, that removes both random and fixed spatial effects and provides the Best Linear Unbiased Estimate (BLUE) of the mean batch canopy size. We apply spatial effect removal to the raw canopy data obtained from all available orthophotos.

Raw and spatial corrected vigor data generated in this project and step-by-step protocol can be found at https://data.4tu.nl/private_datasets/uGa-8eKCK2rKvm72kMt4uJFUzKh6xi_0Hw1rGkH2e-c.

### Choice of the timepoint for potato vigor measurement

Our observations indicate that between 47 and 50 DAP (as indicated in Table S1), all plants have emerged, and their canopies have not yet begun to overlap. During this crucial period, the final selection of dates for vigor measurements was made. In 2019, the last measuring dates were chosen, namely 52 DAP for field M, 48 DAP for field K and 50 DAP for field V. In 2020, the second last measuring dates were chosen, namely 47 DAP for field M, 47 DAP for field K and 49 DAP for field V. It is important to note that these specific date choices do not significantly impact the subsequent prediction modelling or alter the conclusions drawn from the analysis, as the data from the last 2-3 time points are highly correlated (Table S2).

### Sampling of seed tubers for microbiome analysis

In both years, seed tubers were harvested between September and October, stored at 4°C until they were sampled in December or January. One-cm-thick cores were sampled from potato heel ends or eyes using a sterilized Ø 0.6-cm metal corer. Cores from 50 seed tubers were pooled into a single sample per compartment per seedlot for each biological replicate. For each of the 6 varieties, we sampled 4 replicates for each of 10 seedlots in year 1 (240 samples per compartment) and 2 replicates of 30 seedlots in year 2 (360 samples per compartment). In total, 30,000 tubers (12,000 in year 1 and 18,000 in year 2) were sampled to access the microbial composition of different tuber compartments, resulting in 600 samples per compartment. These samples were snap frozen in liquid N_2,_ freeze-dried and stored in 50-mL Falcon tubes at −20 °C until further processing.

### Sample grinding

To efficiently grind the samples in a high-throughput manner, the freeze-dried samples in 50-mL Falcon tubes were amended with four 5-mm sterile metal beads per tube and placed in a custom-made wooden adapter in a paint shaker machine (SK550 1.1 heavy-duty paint shaker; Fast & Fluid, Sassenheim, Netherlands) followed by vigorous shaking for 9 min on maximum intensity (indication of rpm). Freeze-dried sprout samples in 2-mL Eppendorf tubes were grinded with one 5-mm sterile metal bead per tube with a TissueLyser II (Qiagen, Germany) at 30 Hz for 1 min.

### DNA isolation, library preparation and sequencing

Genomic DNA was isolated from ±75 mg eye or heel end powder per sample using a Qiagen PowerSoil KF kit (Qiagen, Germany). The KingFisher™ Flex Purification System machine was used for high-throughput DNA isolation. DNA was quantified using a Qubit® Flex Fluorometer with the Qubit dsDNA BR Assay Kit (Invitrogen, Waltham, MA, USA) and normalized to a concentration of 5 ng/µl. The resulting DNA samples were then stored at −20 °C.

Bacterial 16S ribosomal RNA (rRNA) genes within the V3–V4 hypervariable regions were amplified using 2.5 µL DNA template, 12.5 µL KAPA HiFi HotStart ReadyMix (Roche Sequencing Solutions, Pleasanton, USA), 2 µM primers B341F ( 5’-TCGTCGGCAGCGTCAGATGTGTATAAGAGACAGCCTACGGGNGGCWGCAG-3’) and B806R (5’-GTCTCGTGGGCTCGGAGATGTGTATAAGAGACAGGACTACHVGGGTATCTAATCC-3’) ^43^ with Illumina adapter sequences in combination with 2.5 µM blocking primers mPNA (5’-GGCAAGTGTTCTTCGGA-3’) and pPNA (5’-GGCTCAACCCTGGACAG-3’) in 25 µL reactions. Blocking primers were used to avoid the amplification of mitochondrial (mPNA) or plastidial (pPNA) RNA from the plant host ^44^. Cycling conditions for 16S rRNA were (1) 95 °C for 3 min; (2) 95 °C × 30 s, 75 °C × 10 s, 55 °C × 30 s, 72 °C × 30 s, repeated 24 times; (3) 72 °C × 5 min; (4) hold at 10 °C.

Fungal internal transcribed spacer 2 (ITS2) DNA was amplified using 2.5 µL DNA template, 12.5 µL KAPA HiFi HotStart ReadyMix, 2 µM primers fITS7(5’-TCGTCGGCAGCGTCAGATGTGTATAAGAGACAGGTGARTCATCGAATCTTTG-3’) and ITS4-Rev (5’-GTCTCGTGGGCTCGGAGATGTGTATAAGAGACAGTCCTCCGCTTATTGATATGC-3’) with Illumina adapter sequences in combination with 2 µM blocking primers cl1ITS2-F (5’-CGTCTGCCTGGGTGTCACAAATCGTCGTCC-3’) and clITS2-R (5’-CCTGGTGTCGCTATATGGACTTTGGGTCAT-3’) in 25 µL reactions ^43^. Cycling conditions for ITS2 were (1) 95 °C for 3 min; (2) 95 °C × 30 s, 55 °C × 30 s, 72 °C × 30 s, repeated 9 times; (3) 72 °C × 5 min; (4) hold at 10 °C.

For both PCR reactions, DNA was cleaned using the KingFisher™ Flex Purification System. Twenty µL of vortexed AMPure XP Beads (Beckman Coulter, Brea, USA) were added to 25 µL of PCR product in a KingFisher™ 96 deep-well plate. Beads with adjoined DNA were washed by subsequent transfer to 3 KingFisher™ 96 deep-well plates with 80% ethanol and DNA were then eluted in 30 µL C6 elution buffer from the Qiagen Powersoil KF kit.

Index PCR reactions were performed using standard Illumina i7 (N701-N712) index primers for columns and Illumina i5 (N501-N508) index primers for rows of each plate. Five µL DNA sample was added to a mix of 2.5 µL 2 µM index primer, 12.5 µL KAPA HiFi HotStart ReadyMix and 5 µL Milli-Q H2O. Cycling conditions for index PCRs were (1) 95 °C for 3 min; (2) 95 °C × 30 s, 55 °C × 30 s, 72 °C × 30 s, repeated 9 times for 16S or 24 times for ITS2; (3) 72 °C × 5 min; (4) hold at 10 °C. After the index PCR, DNA was cleaned using the abovementioned cleaning protocol. DNA concentrations of all PCR products were measured using a Qubit® Flex Fluorometer with the Qubit dsDNA BR Assay Kit (Invitrogen, Waltham, MA, USA) and normalised to 2 ng/µL, after which the samples were pooled and sent for Illumina V3 2×300 bp MiSeq sequencing at USEQ (Utrecht, the Netherlands). Step-by-step protocols for DNA isolation and library preparations are available at https://doi.org/10.5281/zenodo.10955437.

### Microbial community analysis

Both 16S and ITS2 rDNA raw sequencing reads were denoised, joined, delineated into amplicon sequence variants (ASVs), and assigned taxonomy in the Qiime2 (v.2019.7) environment ^51^. Datasets were demultiplexed and then filtered using the DADA2 pipeline^52^. ASVs with less than 30 reads or present in less than 3 samples across all samples within a dataset were removed to minimize potential errors in sequencing. The representative sequences were subsequently taxonomically classified using a classifier trained with the 99% OTU threshold SILVA database ^53^ for bacteria and UNITE reference database (v.8.0) for fungi ^54^. For bacteria, we removed remaining 16S reads annotated as mitochondria or chloroplasts and kept only reads assigned to Bacteria. For fungi, we removed remaining ITS reads assigned as *Viridiplantae* and *Protista* and kept only reads assigned to Fungi. The datasets from seed tubers samples were rarefied to respectively 8000 bacterial and 4000 fungal reads per sample. Bray-Curtis dissimilarity matrices were created in QIIME2 and visualized in R using the Qiime2R and *ggplot2* package. Permutational multivariate analysis of variance (PERMANOVA, 999 permutations) tests were performed using QIIME2 to test the effect of different factors on the microbiome composition.

### Modelling and statistics

RF Regression models were built on various taxonomy ranks including ASV, species, genus, family, order, class, phylum and features generated by HFE ^28^ using the *randomForest* package in R with default parameters ^27, 55^. Models were developed independently for both eye or heel end compartment using bacterial, fungal and a combination of both bacterial and fungal datasets. To assess the out-of-bag (OOB) error, the predicted values of the input data were generated using out-of-bag samples. This method involves utilizing data points that were not included in the bootstrap sample used to train a particular model.

We trained the models on the microbiome samples of year 2 and CSA in trial field M in the same year (training set). We tested the model by correlating the predicted CSA to the observed CSA in the two other trial fields of the same year (within-year testing set). We then tested the RF model further with the tuber microbiomes from year 1 (across-year testing set) originating from a completely distinct collection of seedlots.

The seedlots were divided into three classes of high (H), medium (M), and low (L) vigor by determining the 3-quantiles of the predicted and observed vigor. The 3-quantiles are values that divide a sample vector into three portions of equal probability, in our case, when the probability distribution of vigor is unknown, these are values that divide a given vector into portions of the same size, e.g. the median of a vector is its 2-quantile. Samples have low vigor if their measured or predicted vigor measure is smaller than the first 3-quantile, medium if it is between the first and second 3-quantile, high if it is greater than the second 3-quantile.

To measure the performance of the classification model, we compute the class specific precision measure for a class Ci as:

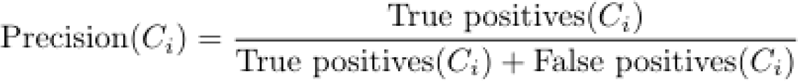

i.e. the ratio of correctly classified samples in Ci over the total number of samples classified as Ci. Note that in terms of conditional probabilities this is equivalent to:

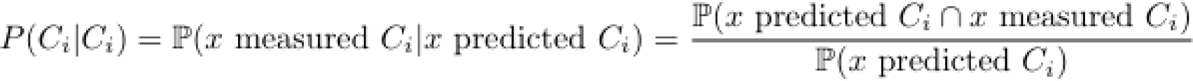

Intuitively, a good classification model will have values close to one on the diagonal and low values off diagonal, indicating that the probability of a sample to be misclassified is much lower than the probability of being classified correctly. We define the “quality confidence” of the prediction as one minus the probability of the sample being misclassified as the opposite extremum. Whether the quality confidence is significantly higher than random chance was indicated by pseudo-*P* generated with 1000 times simulation of random classification. (All codes will be made available)

### Identification of keystone microbes

The importance of each ASV was extracted from the model using the *importance()* function in the *randomForest* package^27^. Briefly, this importance metric represents the change in the error of the model after permuting the counts of each of the ASVs and is therefore a measure of the contribution of each separate ASV to the model. This change is calculated for different permutations of the OOB data, averaged across trees, and normalized by the standard deviation of all values. The top 1% and 5% of these values were used to obtain the most strongly contributing ASVs. Partial dependence is defined as the individual effect of each ASV to the model prediction and was calculated using the *hstats* package^56^ in R for the top 1% most contributing ASVs.

## Supporting information

Song et al Supplementary figures and tables

## Acknowledgements

We gratefully acknowledge the valuable contributions of the Royal HZPC Group and Averis Seeds B.V. in providing the seed tuber material and supporting the field trials. Their collaboration was essential to the successful execution of this research. Special thanks are extended to Falko Hofstra and Martzen ten Klooster from HZPC Holding B.V. for their contribution to the sample collection and Johan Hopman from Averis seeds B.V for his valuable advice. We also thank Casper Jongekrijg and Ellen Manders and Emma de Kloe for excellent technical assistance in the laboratory. Additionally, we acknowledge the funding support received from Europees Landbouwfonds voor Plattelandsontwikkeling (ELFPO) on the “Flight-to-vitality” project. This work was also partly supported by the Dutch Research Council (NWO) through the Gravitation program MiCRop (grant no. 024.004.014) and through project “Sequence-based POTato Microbiome tools for microbiome-optimized potatoes” (project no. 19769).

## Data availability

The data that support the findings of this study are available from the corresponding author upon reasonable request. Moreover, the raw sequence data generated of this study are available at https://dataview.ncbi.nlm.nih.gov/object/PRJNA1091851?reviewer=evm2u22thjvhe2d9bgut1ecrkt

## Code availability

Raw and spatial corrected vigor data generated in this project and step-by-step protocol are available at https://data.4tu.nl/private_datasets/uGa-8eKCK2rKvm72kMt4uJFUzKh6xi_0Hw1rGkH2e-c.

## Authors and Affiliations

Yang Song, Juan J. Sanchez Gil, Ronnie de Jonge, Peter G.H. de Rooij, David Kakembo, Peter A.H.M. Bakker, Corné M.J. Pieterse, Roeland L. Berendsen **Plant-Microbe Interactions, Institute of Environmental Biology, Department of Biology, Science4Life, Utrecht University, Padualaan 8, 3584 CH Utrecht, the Netherlands.**

Eliza Atza, Neil V. Budko, **Numerical Analysis, Delft Institute of Applied Mathematics, Delft University of Technology, Mekelweg 4, 2628 CD Delft, the Netherlands.**

Doretta Akkermans, **HZPC research B.V. Roptawei 4, 9123 JB Mitselwier, the Netherlands**

## Contributions

YS, DA, EA, NVB, RJ, PAHMB, CMJP and RLB designed the experiments and approach. YS, EA, JJSG, NVB, PAHMB, CMJP, and RLB wrote the manuscript. DA coordinated sampling collection and experimental field trials. YS and PGHR carried out molecular analysis. EA and NVB performed quantitative analysis of drone images and CSA data. YS, EA, JJSG, RJ, DK performed data analysis.

## Ethics declarations

The authors declare that this study received funding from HZPC Research B.V. and Averis Seeds B.V. The funder had the following involvement in the study: study design, sample collection, and the decision to submit it for publication. DA is a current employee of HZPC research B.V.

