## Supplementary material for "Seed tuber microbiome is a predictor of next-season potato vigor": Song et al Supplementary figures and tables

**The following Supporting Information is available for this article:**

Supplementary Figures

Figure S1. **Experimental design over 2 years.**

Figure S2. **From field images to potato vigor data.**

Figure S3. **Development of CSA per trial field and year.**

Figure S4. **Seedlot variation in CSA of the Festien variety in field M in year 1.**

Figure S5. **Effects of each potato seedlot on potato vigor for all varieties and both years.**

Figure S6. **Microbiomes of replicate samples of the same seedlot cluster together in year 1.**

Figure S7. **Microbiomes of replicate samples of the same seedlot cluster together in year 2.**

Figure S8. **Seed tuber microbiomes in heel end compartments.**

Figure S9. **Scatter plots illustrating the Pearson correlation between the predicted and observed potato vigor in all fields and varieties.**

Figure S10. **General assessment of the relationship between the top 1% contributors to potato vigor.**

Figure S11. **Heatmaps showing Spearman correlations of each of the top 1% contributing ASVs to potato vigor, their prevalence, and median abundance across samples.**

Supplementary Tables

Table S1. **Summary of the canopy surface area (CSA) measurements by drone imaging of the three test fields (M, V, and K) in year 1 and year 2.**

Table S2. **Correlations in potato vigor between trial fields for all varieties or each variety.**

Table S3. **List of seedlots selected for the microbiome analysis in year 1 and year 2.**

Table S4. **Performance of RF models trained with tuber heel end microbiome at distinct taxonomic ranks.**

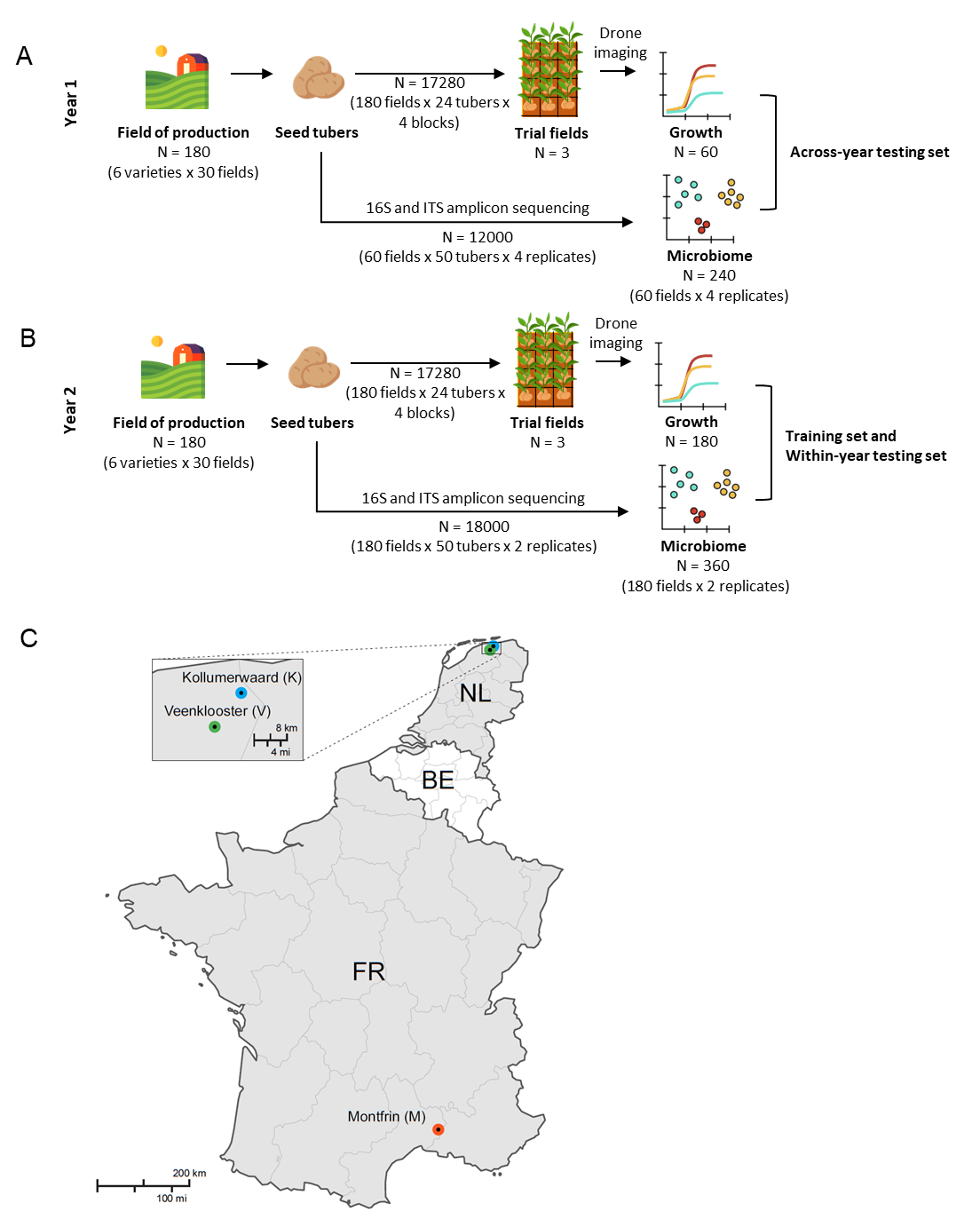
Figure S1. **Experimental design over 2 years.** **A-B** We collected seed tubers of 6 potato varieties from 30 fields per variety (180 fields in total) in the Netherlands in the autumn of 2018 (year 1) and 2019 (year 2). Tubers from these 180 fields per year were stored over winter and the tubers were planted in each of 3 trial fields in the next spring. **C** In both years, the trial fields were located near Montfrin (M) in France and near Kollumerwaard (K) and Veenklooster (V) in the Netherlands. In each of the trial fields, the seed tubers were planted in randomized block design with 4 replicate blocks of 24 tubers. We monitored the growth and development of the plants that emerged from these seed tubers using aerial images of the complete field with a drone-mounted camera. Of the 180 seedlots of year 1, we selected 60 seedlots from which we took 4 replicate samples for microbiome analysis. In year 2, the microbiomes were analyzed of 2 replicate samples of all 180 seedlots.

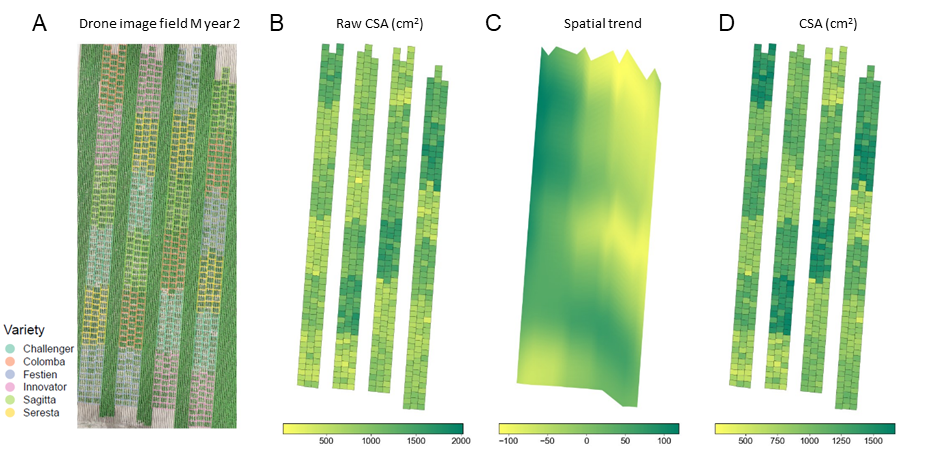
 Figure S2. **From field images to potato vigor data.** **A** Exemplary ortho image of trial field M in year 2 obtained with drone-mounted camera. Plot boundaries of each seedlot are displayed in variety-specific colors. **B** Overview of raw canopy surface area (raw CSA) per plot in the trial field displayed as a heatmap. **C** Spatial trend of the trial field recovered with the SpaTS package and displayed as a heatmap. **D** Overview of spatially corrected raw CSA in the trial field as a heatmap. Average corrected seedlot CSA is shown in all replicate plots of the four seedlots.

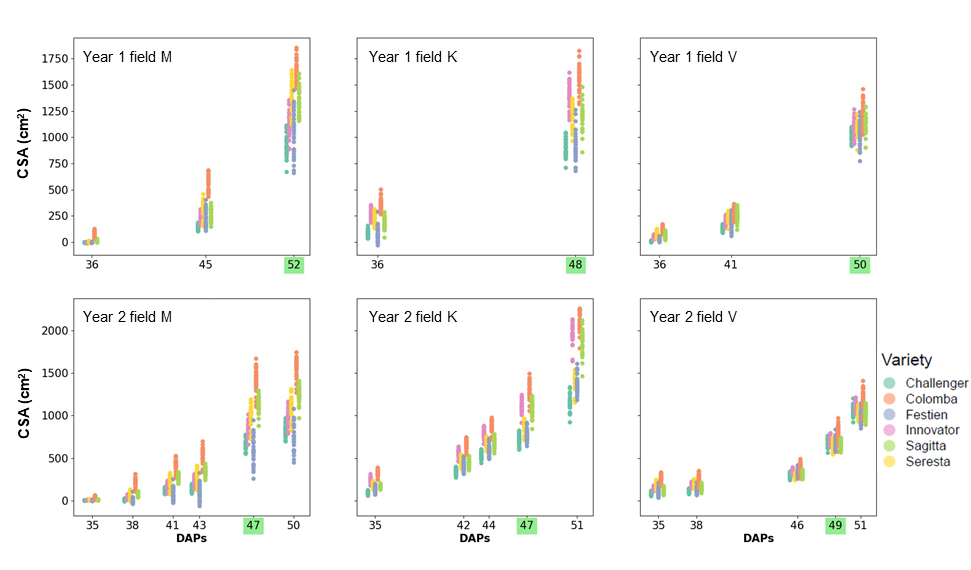
Figure S3. **Development of CSA per trial field and year.** Each symbol represents the average CSA of 4 replicate plots of a seedlot after correction for field spatial effects. In each plot, *x*-axis represents days after planting (DAP), *y*-axis represents CSA (cm^2^). Each variety is shown in a different color as indicated in the legend. The dates chosen for the regression studies are highlighted in green.

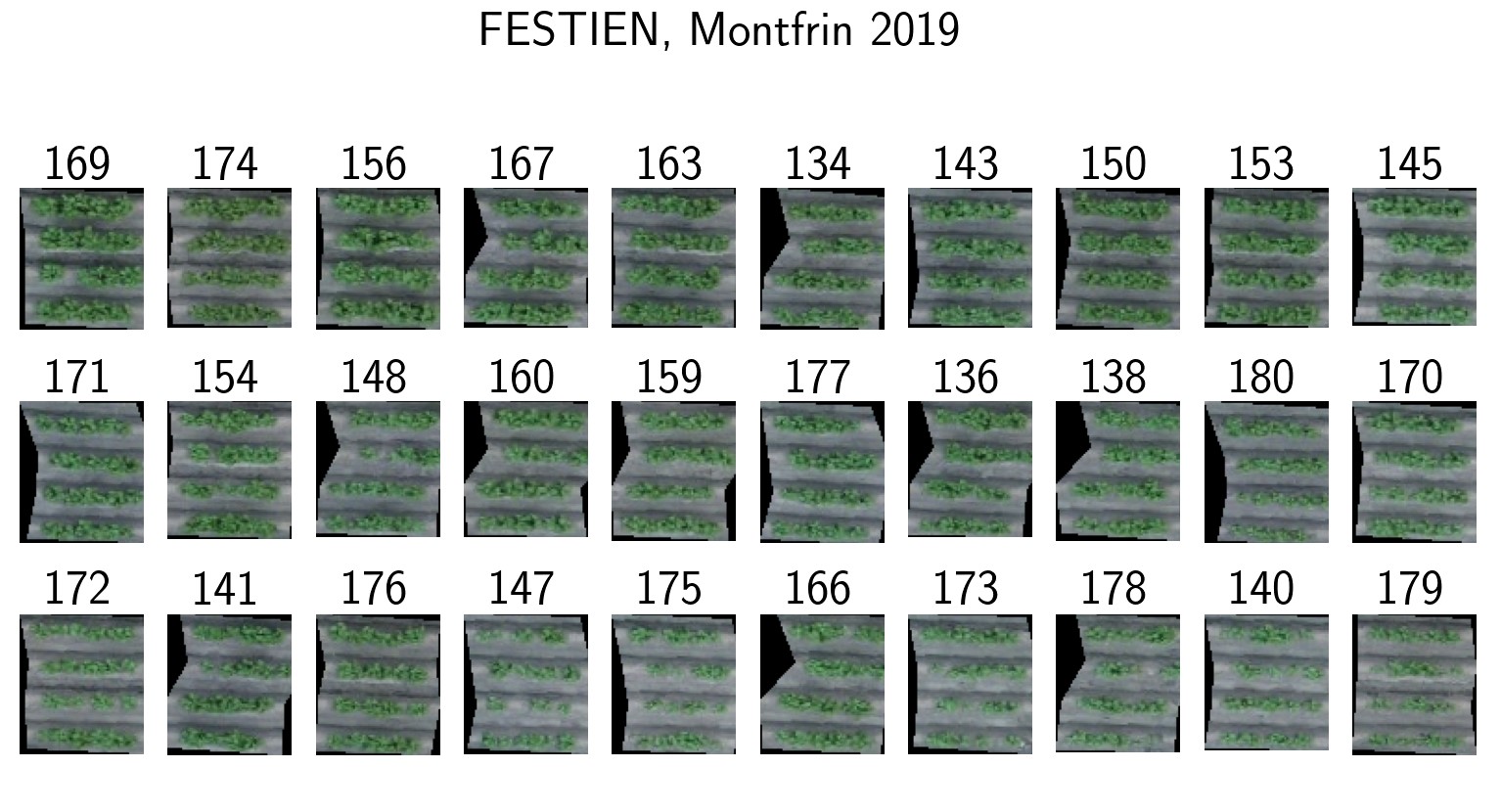
Figure S4. **Seedlot variation in CSA of the Festien variety in field M in year 1.** Drone images of a representative plot of the 4 replicate plots of each seedlot in the field. Numbers labeled on top of each plot indicate the seedlot number of the plot.

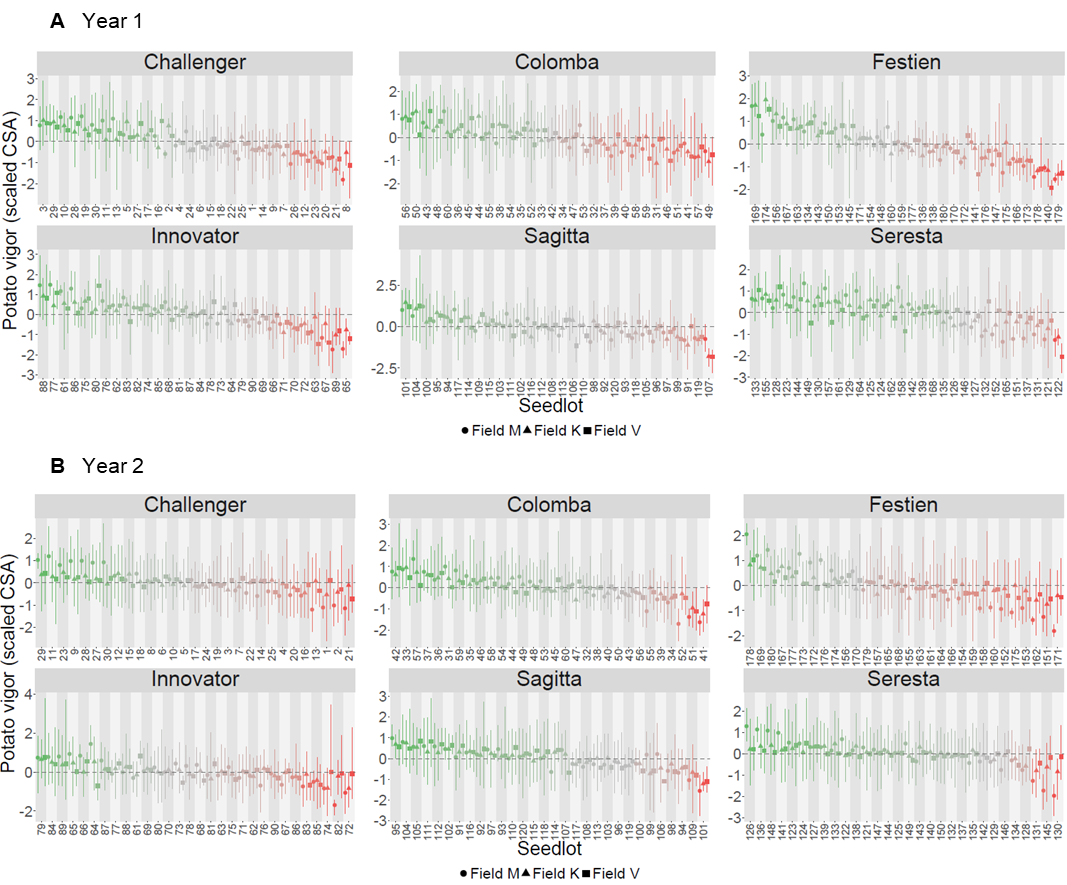
Figure S5. **Effects of each potato seedlot on potato vigor for all varieties and both years. A** Scaled CSA for each of the 6 varieties and each of the 30 seedlots per variety in Field M, K and V in year 1. **B** Scaled CSA for each of the 6 varieties and each of the 30 seedlots per variety in Field M, K and V in year 2. Error bars signify the minimum and maximum values for a given seed lot per trial field. The CSA in each trial field, as estimated by the SpaTS package, is indicated with the field corresponding marker (see legend).

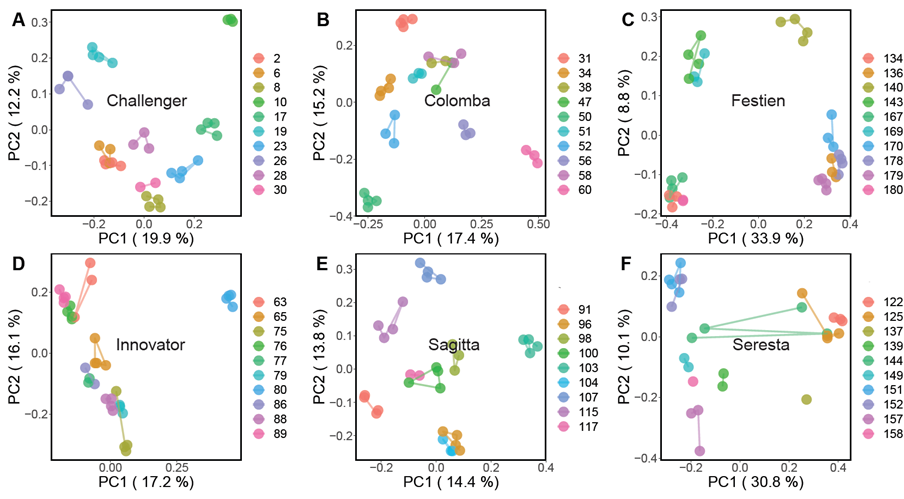
Figure S6. **Microbiomes of replicate samples of the same seedlot cluster together in year 1.** PCoA ordination plot based on Bray-Curtis dissimilarities of bacterial communities of seed tuber eye compartment from year 1. Variety names are indicated in each panel. Each data point represents a replicate of one seedlot. Different colors represent different seedlots of a variety.

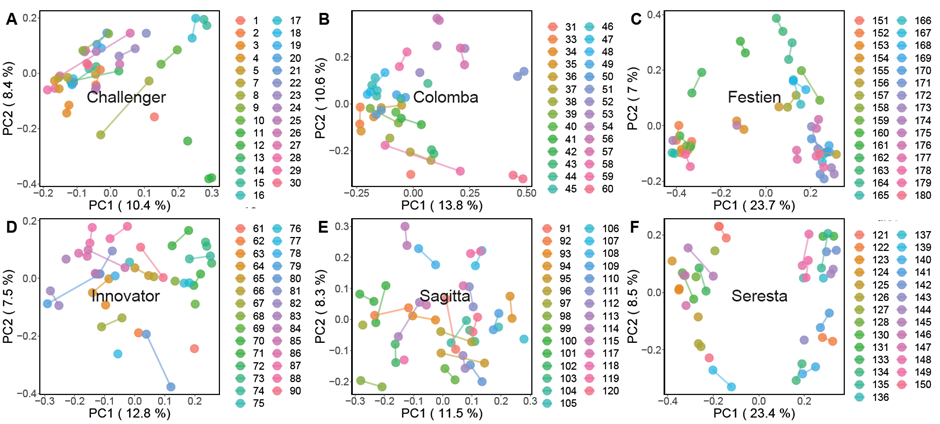
 Figure S7. **Microbiomes of replicate samples of the same seedlot cluster together in year 2.** PCoA ordination plot based on Bray-Curtis dissimilarities of bacterial communities of seed tuber eye compartment from year 2. Variety names are indicated in each panel. Each data point represents a replicate of one seedlot. Different colors represent different seedlots of a variety.

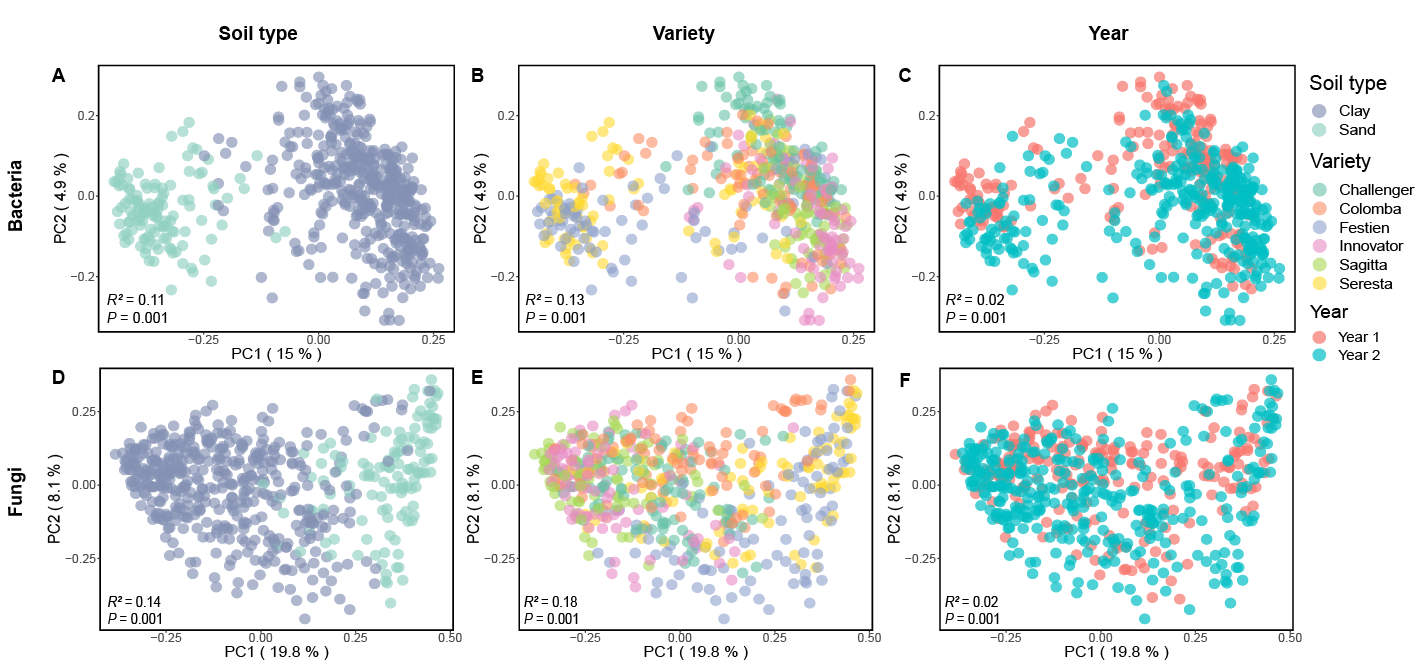
Figure S8. **Seed tuber microbiomes in heel end compartments.** Principle coordinate analysis (PCoA) ordination plot based on Bray-Curtis dissimilarities of bacterial (**A-C**) and fungal (**D-F**) microbiomes. Symbols are colored by soil type (**A,D**), variety (**B,E**) and year (**C,F**) as indicated in the legend. Each data point represents a single replicate of a seedlot. Four replicate samples were analyzed for each of 60 seedlots in year 1 and two replicate samples for each of 180 seedlots in year 2.

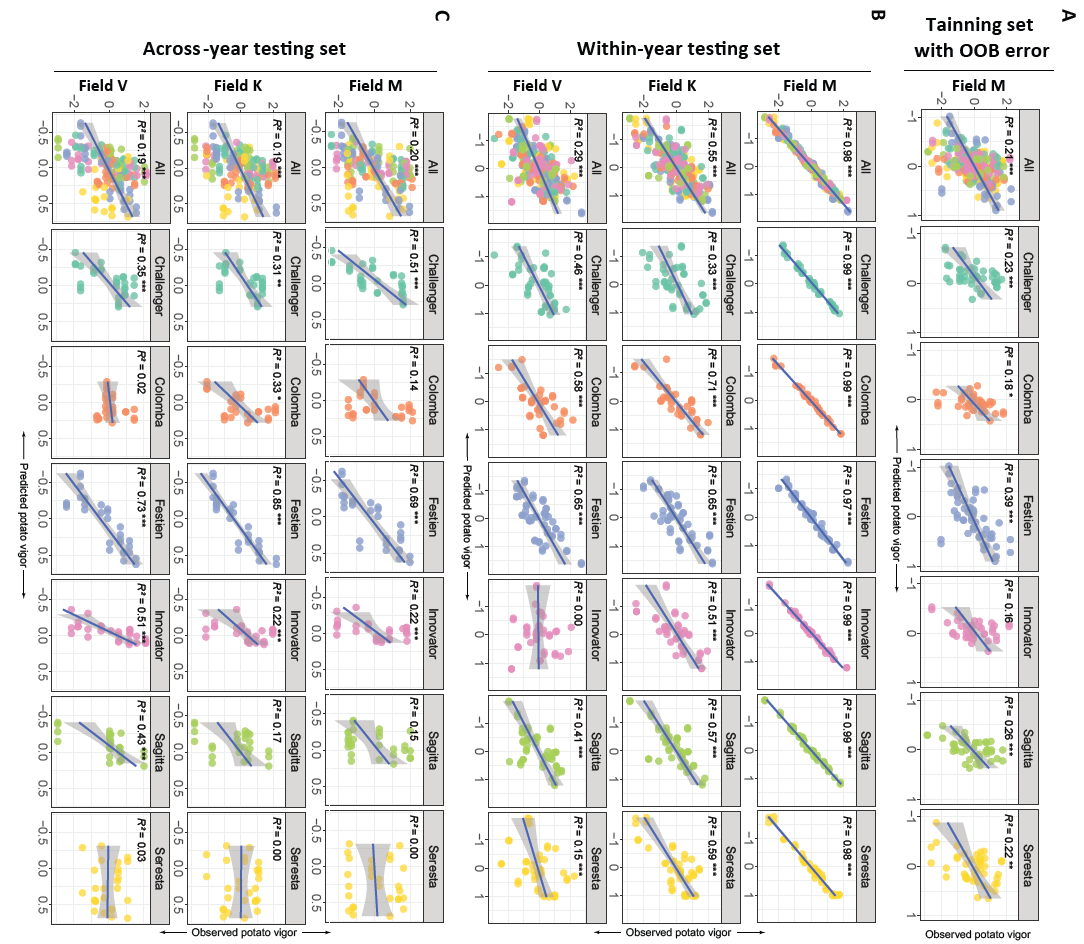
 Figure S9. **Scatter plots illustrating the Pearson correlation between the predicted and observed potato vigor in all fields and varieties.** In all panels, values on the x-axis are predicted by a random forest model trained on microbiome data from year 2 and CSA from field M. The predicted potato vigor is based on the same microbiome data as was used for training the model (within-year testing set) or based on microbiome data from year 1 to which the model was naïve (across-year testing set). The 6 varieties are represented by different colors. Each symbol represents a prediction microbiome based on 1 eye compartment sample. Predicted and observed vigor are indicated by scaled CSA, which is scaled to the variety average in each trial field.

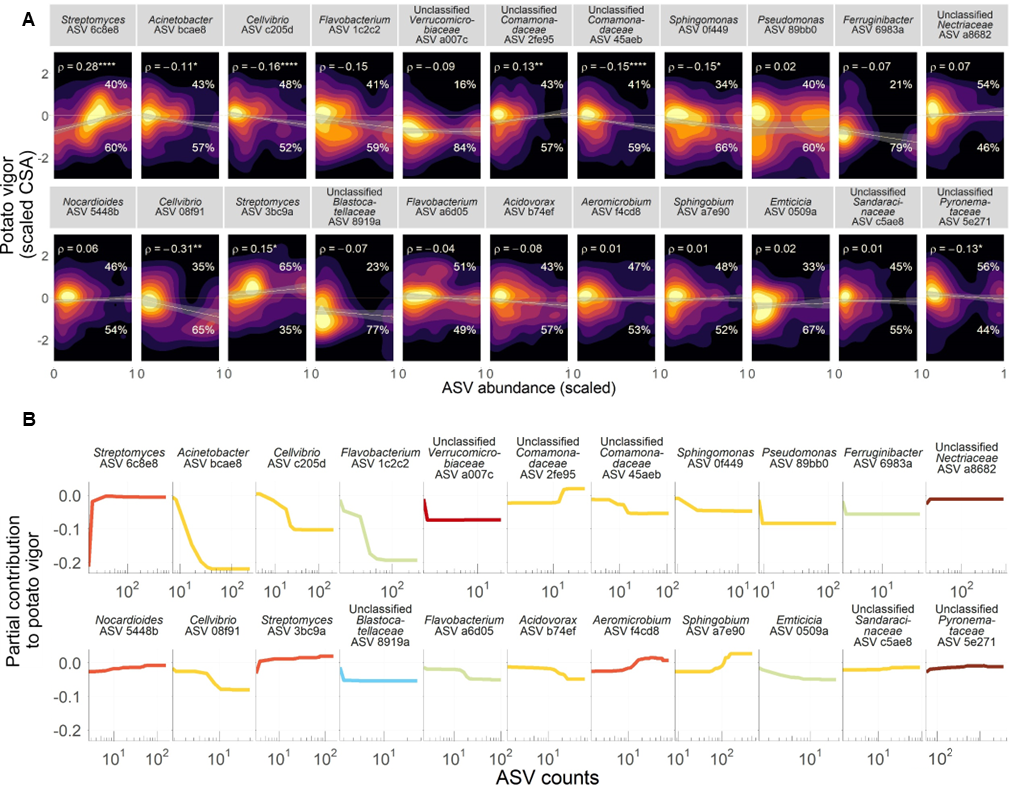

Figure S10. **General assessment of the relationship between the top 1% contributors to potato vigor**. **A** Bidimensional density plots showing scaled CSA values and normalized abundance of each of the top 1% contributing ASVs. ASV abundance is rescaled between 0 and 1 with respect to their minimum and maximum in order to show one single scale across ASVs. The clearest colors indicate areas that accumulate most of the data, and dark colors the areas where no data or few points are found. The line was fitted with a robust regression to outliers computed with the *rlm()* functions in the *MASS* R package, and the *ρ* values indicate Spearman’s *ρ* together with the significance level shown by asterisks (**P*<0.05; ***P*<0.01; ****P*<0.001; *****P*<0.0001). The percentages above and below the 0-line indicate the number of ASV occurrences in sample with vigor above and below the mean, respectively. **B** Partial contribution plots for the top 1% ASVs most predictive to potato vigor (scaled CSA) according to the RF model.

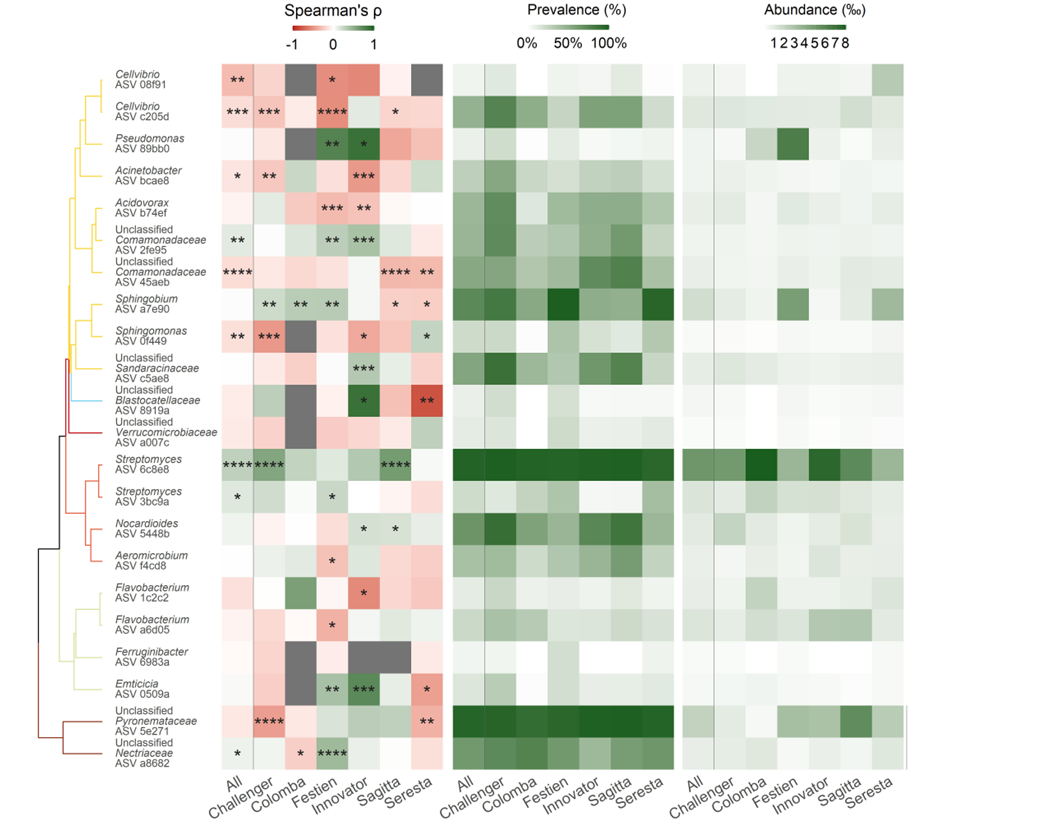

Figure S11. **Heatmaps showing Spearman correlations of each of the top 1% contributing ASVs to potato vigor, their prevalence, and median abundance across samples.** The first column in every heatmap shows the computed value including all the data regardless of plant variety, and the rest of columns display those values calculated for individual potato varieties. The abundance of the fungal ASVs were shown as 1/10 of the original value to fit in the color scale. The significance level of Spearman’s *ρ* are shown by asterisks (**P*<0.05; ***P*<0.01; ****P*<0.001; *****P*<0.0001).

Table S1. **Summary of the canopy surface area (CSA) measurements by drone imaging of the three test fields (M, V, and K) in year 1 and year 2.** The column ‘Date’ shows the dates (year-month-day) of the measurement. The column ‘DAP’ shows the time of the drone image in days after planting (DAP). The column ‘zero ridges (%)’ gives the number of ridges (each plot has four ridges) with no measurable canopy and their fraction among all ridges in the field.

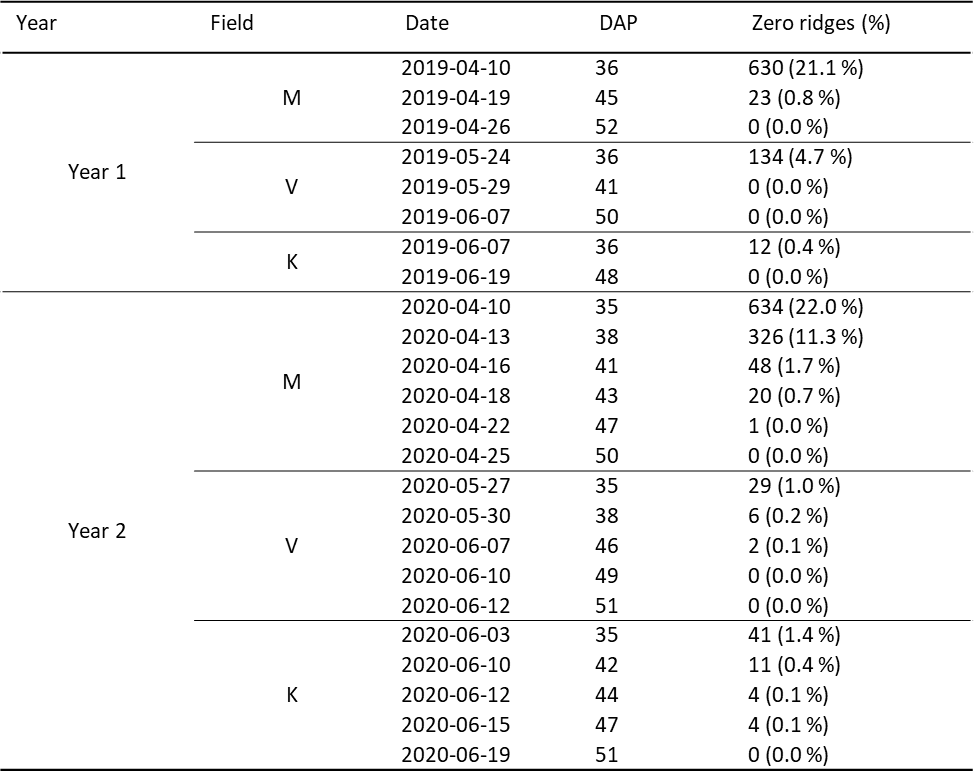

Table S2. **Correlations in potato vigor between trial fields for all varieties or each variety.** * indicates *P* lower than 0.05, ** indicates *P* lower than 0.01 and *** indicates *P* lower than 0.001.

| Year | Variety | M-V | M-K | V-K |
| --- | --- | --- | --- | --- |
| Year 1 | All | 0.52*** | 0.72*** | 0.64*** |
|  | Challenger | 0.77*** | 0.63*** | 0.76*** |
|  | Colomba | 0.41* | 0.66*** | 0.59*** |
|  | Festien | 0.77*** | 0.77*** | 0.82*** |
|  | Innovator | 0.63*** | 0.86*** | 0.61*** |
|  | Sagitta | 0.33 | 0.57** | 0.64*** |
|  | Seresta | 0.21 | 0.81*** | 0.42* |
| Year 2 | All | 0.54*** | 0.74*** | 0.52*** |
|  | Challenger | 0.64*** | 0.63*** | 0.26 |
|  | Colomba | 0.71*** | 0.79*** | 0.73*** |
|  | Festien | 0.74*** | 0.76*** | 0.66*** |
|  | Innovator | 0.11 | 0.71*** | 0.23 |
|  | Sagitta | 0.64*** | 0.78*** | 0.81*** |
|  | Seresta | 0.43* | 0.78*** | 0.44* |

| Variety | Seedlots year 1 | Seedlots year 2 |
| --- | --- | --- |
| Challenger | 2, 6, 8, 10, 17, 19, 23, 26, 28, 30 | 1-30 |
| Colomba | 31, 34, 38, 47, 50, 51, 52, 56, 58, 60 | 31-60 |
| Festien | 214, 216, 220, 223, 247, 249, 250, 258, 259, 260 | 61-90 |
| Innovator | 63, 65, 75, 76, 77, 79, 80, 86, 88, 89 | 91-120 |
| Sagitta | 91, 96, 98, 100, 103, 104, 107, 115, 117 | 121-150 |
| Seresta | 202, 205, 217, 219, 224, 229, 231, 232, 237, 238 | 151-180 |

Table S3. **List of seedlots selected for the microbiome analysis in year 1 and year 2.**

Table S4. **Performance of RF models trained with tuber heel end microbiome at distinct taxonomic ranks.** RF models trained on potato vigor and tuber microbiome data from field M in year 2 were tested on vitality data from field M (training set), field K and V in year 2 (within-year testing set) and then tested on data from field M, K and V in year 1 (across year testing set). Numbers represent the *R*^2^ of a model, where a higher value indicates superior performance. Numbers in black represent significant correlations where *P* < 0.05, gray represent unsignificant corrections. The row of “OOB” indicates the out-of-bag (OOB) performance of the models in the training set.

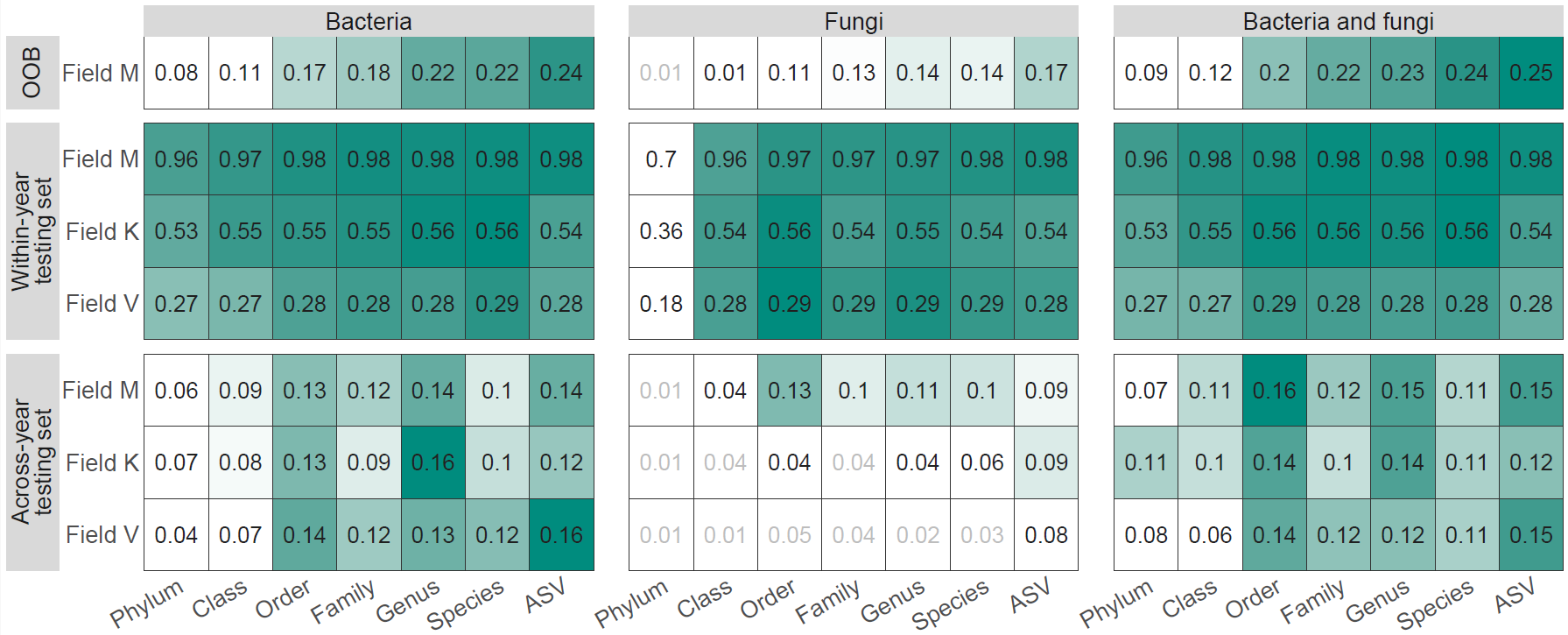
